# Inference and validation of a dynamic gene regulatory network influencing cell fate during Duchenne muscular dystrophy initiation

**DOI:** 10.1101/2025.11.19.689189

**Authors:** Elsa G Guillot, Quentin G Fort, Elise Mozin, Bérangère Robert, Matteo Bouvier, Cynthia Fourgeux, Paul Simon, Sara Munoz, Matthew Childers, Jeremie Poschmann, Oumeya Adjali, David L Mack, Arnaud Bonnaffoux, Jean-Baptiste Dupont

## Abstract

The onset of gene regulatory disturbances leading to developmental disorders remains poorly understood, particularly when symptoms begin after birth. Using induced pluripotent stem cells, single-cell transcriptomics and computational modeling, we determined the gene expression waves guiding early myogenesis and how they are affected by a disease-causing mutation. We uncovered a dynamic gene regulatory network organized in two kernels that function as a toggle-switch controlling stochastic cell fate decisions. In cells derived from patients with Duchenne muscular dystrophy, these waves of regulation are less pronounced and the switch remains in the OFF state with a higher probability, resulting in convergence into a restricted cell fate. Experimental reactivation of the switch through RNA interference led to a rescued transcriptomic state and to a refined network topology. This work uncovers a dynamic regulatory mechanism underlying early cell fate divergence and provides a general framework to infer and validate dynamic gene regulatory networks involved in disease initiation.

## Introduction

Duchenne Muscular Dystrophy (DMD) is a genetic disease caused by mutations in the *DMD* gene encoding the dystrophin protein. Diagnosis occurs when delays appear in the acquisition of early motor milestones in young patients. However, alterations of the muscle histology have been described more than fifty years ago in asymptomatic newborns and in fetuses at risk of developing DMD^1–3^. To better understand the dynamics of DMD initiation, induced pluripotent stem cells (iPSCs) derived from patients are of major interest as they can recapitulate the early steps of human embryonic development and disease in a dish. Initial studies investigated the impact of the DMD mutations on the later stages of iPSC differentiation leading to terminally differentiated myotubes. They showed that the absence of dystrophin correlates with an impaired ability to form myogenic progenitors and an increased proportion of branched myotubes^4^. At the histological level, the number of nuclei per myotube was found significantly reduced, which indicates limited differentiation capabilities^5^. In light of these findings, we investigated the impact of dystrophin mutations on earlier developmental stages, including the formation and differentiation of paraxial mesoderm and somite progenitors from iPSCs^6,7^. Using bulk RNA-Sequencing (RNA-Seq), we highlighted a striking transcriptome dysregulation in DMD cells at the somite stage^6^. At single-cell resolution (scRNAseq), we established the differentiation trajectory and observed that most – but not all – DMD cells bifurcate to a pathological state characterized by a marked dysregulation of cell junction genes and an impaired ability to form epithelial cell clusters^7^. This result is of particular interest, since it suggests that dystrophin deficiency could impact early development, particularly somite formation and patterning. However, the exact regulation mechanisms leading to the pathological state remains to be investigated. In this study, our objective is to decipher the gene regulatory network (GRN) driving cell fate choice at the bifurcation point in DMD cells, and to validate it experimentally.

Seminal work on single-cell trajectory analysis highlighted the non-determinism of cell fate during DMD initiation^7^. While each branch of the trajectory contained 86% of cells from a single iPSC line (either Healthy or DMD), we still observed 14% of cells from the other genetic background. This indicates that the choice between these two genetic states is probabilistic and does not strictly depend on the presence or absence of the DMD mutation. Here, we raise the hypothesis that these probabilities are determined by a GRN with a toggle switch topology^8,9^ that is perturbed - directly or indirectly - by the DMD mutation. Toggle switch patterns are known to regulate biological processes in vivo such as hematopoiesis^10^ or neural development^11^, and they have also been reproduced in silico in synthetic systems^12–14^. A toggle switch is composed of two kernels, i.e. groups of genes interacting in a specific pattern: auto-activation loops between genes of the same kernel, and cross-inhibition between kernels. This creates two stable “attractors” (i.e. cell fates) to which cells converge in a mutually exclusive manner, with a ratio between attractors depending on toggle switch competitive interactions balance. When considering a differentiation process at single-cell resolution, this leads to a bifurcation in the trajectory. In our scenario, the DMD mutation does not drastically change the network topology, which would have created a new cellular state, but rather unbalances the attractors and modifies the strength of specific interactions within the GRN. This assumption implies to adopt a dynamic, causal, mechanistic and probabilistic GRN inference method. An increasing number of inference algorithms are being developed based on the analysis of single cell OMICS data^15,16^. Some purely rely on descriptive statistics approaches and thus are limited to correlations. Others assume simple linear or binary mechanistic models. Here, we adapted the WASABI pipeline previously developed in our group for the analysis of bifurcations in stochastic trajectories obtained from time series of single-cell transcriptomic data^17^. WASABI is based on a machine learning method built on the principle of wave analysis. A wave refers to the propagation of an external stimulus which initiates a cascade of gene activation or inhibition. Using reverse engineering, gene interactions can be inferred and hierarchized by measuring their quantitative expression waves at the RNA and protein levels, assuming that the expression of a regulatory gene precedes the expression of its regulon. Wave analysis therefore requires a prior feature selection step to decrease the number of genes of interest before GRN inference. This novel retro-engineering approach provides insight into the core regulatory mechanisms involving a limited number of key genes, which can be experimentally validated^18,19^.

In the present study, we adapted the wave analysis approach from WASABI for the inference of the core GRN driving the bifurcation between the Healthy and DMD cell states by exploiting the specific properties of bifurcation trajectories. Importantly, the inferred GRN was then experimentally validated in the iPSC model by perturbation experiments using RNA interference.

## Results

### Identification of the single-cell trajectory of interest during myogenic differentiation of DMD iPSCs

We previously demonstrated that iPSCs derived from a DMD patient and subjected to myogenic differentiation acquire a distinct transcriptomic profile from healthy controls (summarized in Figure 1a-b)^7^. Cells collected at 10 successive time points between the pluripotent and myogenic stages were sequenced and the single-cell trajectory was reconstructed with the Monocle suite. We obtained a branched trajectory with an unbalanced distribution of Healthy and DMD cells on the two daughter branches (Figure 1b). While each branch contained 86% of cells from a single line (either Healthy or DMD), we still observed 14% of cells from the other genetic background. This indicates that the choice between the two genetic states is probabilistic and does not strictly depend on the presence of the DMD mutation. Thus, since phenotypes (Healthy, DMD) and genotypes (wild type, mutant) can be intertwined, we will refer to cells derived from Healthy patients as “wild type” (WT) and to cells derived from DMD patients as “mutant” to avoid any confusion between phenotype and genotype.

**Figure 1:**
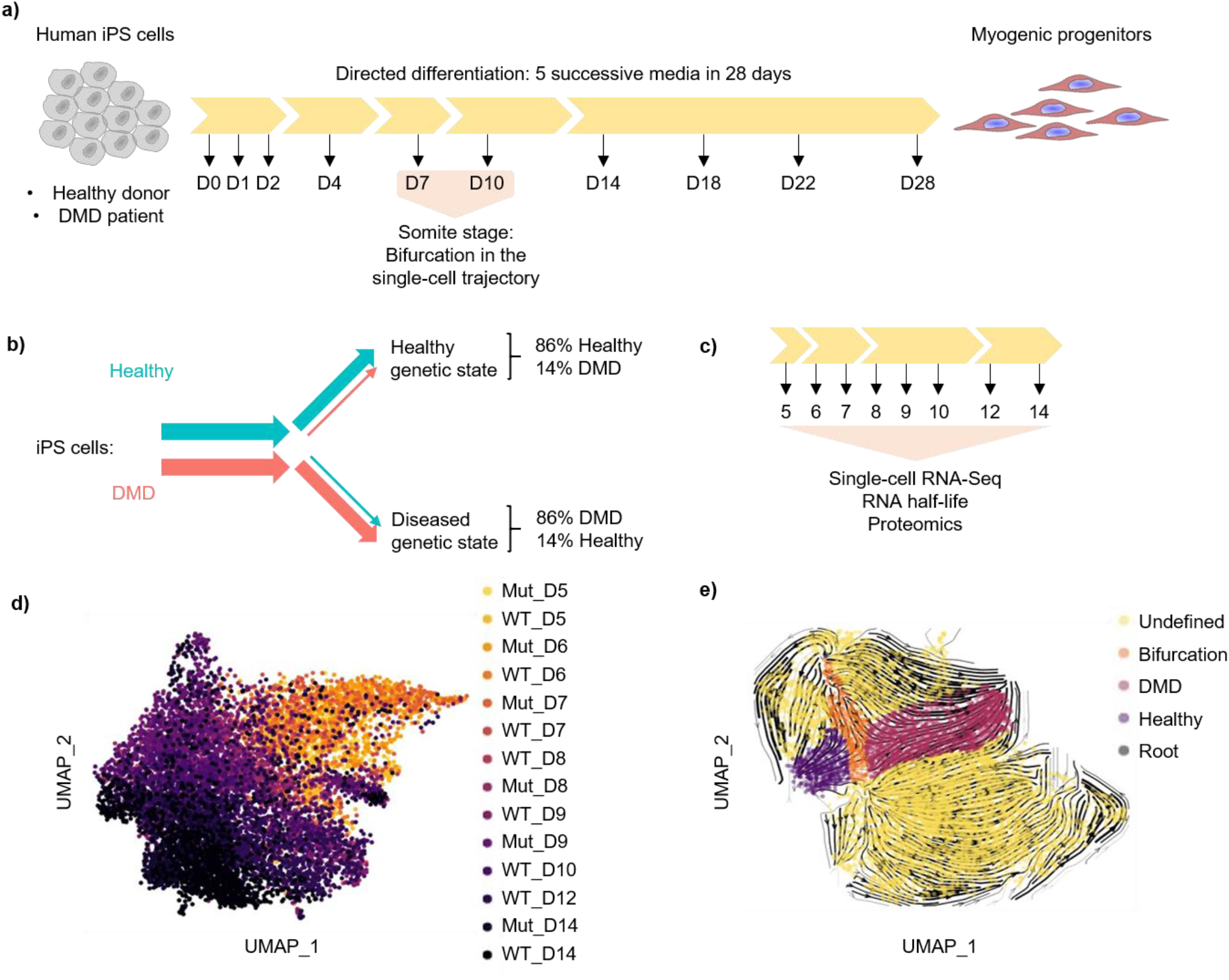
Identification of the bifurcation trajectory of interest. a) iPS cell differentiation into the myogenic lineage with cell collection time points indicated in days (D0 to D28). b) Schematic representation of single-cell trajectory as reconstructed in Mozin 2024 from Healthy and DMD iPS cells using the Monocle pipeline (Trapnell et al., 2017). c) Differentiation pipeline focused on the bifurcation point and molecular analyses performed at eight time points (in days) in the present study. d) UMAP of scRNAseq data with all genes and all cells colored by experimental condition. e) UMAP and RNA velocity of a subset of the original dataset with only cells at D9 to D14 without pluripotent cells considering genes differentially expressed in the bifurcation of interest. The trajectory of interest is labelled with 4 distinct groups: Root, bifurcation and branches Healthy and DMD.

In the present study, we hypothesize that cell fate choice at the bifurcation point is governed by a GRN. The initial step of our inference method is to identify and characterize the genetic trajectory of interest. To gain insight into the molecular events directly downstream of the mutation, we generated more resolutive scRNA-Seq data around the bifurcation point. We subjected WT and mutant iPS cells to directed myogenic differentiation and we used Cellular Indexing of Transcriptomes and Epitopes by sequencing (CITE-Seq)^20^ to barcode cells collected at 8 time points spanning the bifurcation window between Day 5 and Day 14 (Figure 1c). We confirmed the absence of batch effect and first analyzed the scRNA-Seq data considering all genes and all cells with a dimension reduction approach using the Uniform Manifold Approximation and Projection algorithm (Figure 1d, Figures S1-S2)^21^. The RNA velocity data is consistent with the experimental time points (Figures S3-S4) and with the expression of mesoderm development marker genes (Figure S5)^22^. Overall, we observed a large cell cluster expressing somite markers, consistent with the emergence of distinct cell fates which could result in a bifurcation trajectory, as observed in the initial study^7^. However, the trajectory obtained with this new data set and cells collected between Day 5 and Day 14 is more complex and convoluted than the trajectory obtained between Day 0 and Day 28. This result was expected as cell state dispersion is hypothesized to be maximal around the bifurcation.

To deconvolute this blurred trajectory, we performed a series of feature selection steps to subset the most informative cells and genes. We performed a cell density analysis, which revealed overlaps between cells collected at day 5, 6 and 7 (Figure S6) and separation in different sub-clusters between day 9 and 14 (Figure S7). In addition, the analysis of RNA velocity data revealed the existence of 4 final attractors instead of the 2 initially identified as Healthy and DMD (Figure S8). Thus, we focused on the Day 9 to Day 14 window to characterize the dynamics of the attractors, which again detects 4 final attractors and an isolated cluster of cells (Figure S9, S10, Table S3). This isolated cluster shows a high and specific expression of pluripotency marker genes, suggesting that these cells did not undergo differentiation (Figure S11) and should be discarded from our analysis. A novel UMAP was built from RNA velocity data excluding the cells expressing pluripotency markers (Figures S12 and S13). Here again, four attractors were detected, including the two previously identified as Healthy and DMD^7^, which were respectively defined as the terminal Healthy and DMD states in the present study. Interestingly, while WT cells were found in both attractors, mutant cells were found almost exclusively in the DMD attractor. In other words, WT cells have the potential to adopt either the Healthy and the DMD cell fate, while the mutant cells are almost exclusively restricted to the DMD cell fate.

Subsequently, an additional feature selection step was performed to reduce the number of genes considered in the trajectory analysis by keeping only the genes whose expression level varies along the trajectory (Figure S14). A novel UMAP was generated based on this set of genes. As expected, four final attractors were identified, including the two attractors corresponding to the terminal states Healthy and DMD (Figure 1e, S15, S16, S17). The two other attractors show strong expression of genes respectively involved in the cell cycle and neuronal differentiation (Fig S18, S19). This trajectory suggests that between D5 and D14, a significant part of the cells follow the expected myogenic differentiation trajectory and reach either the Healthy or DMD cell fate (Table S4, S5). The remaining cells and their respective trajectories are considered out of interest in the context of the present study, as they represent undifferentiated cells involved in cell division or cells progressing on an aberrant differentiation path, as previously observed with this differentiation protocol^23^. In the next step of our analysis, we focused on the bifurcation trajectory obtained after the successive feature selection steps described above and shown in Figure 1e to infer the GRN governing cell fate decision.

### Inference of the gene regulatory network driving the bifurcation trajectory

We then used the WASABI framework to estimate and compare the waves of gene expression along the pseudotime of the 2 branches detected in the trajectory. WASABI detected 692 waves which were filtered to detect genes with a characteristic toggle switch expression pattern (Figures 2a-c). Ultimately, we obtained 6 core genes meeting these criterias: *ENO1, ID1, MIF, PDGFRA, PRTG* and *PTN* (Figure 2d-e, Figures S21-S22). The expression pattern of these genes satisfies the necessary conditions described in the material and methods section: (i) differential wave patterns between the healthy and DMD trajectories (Figure 2d, Figures S23, S24, S25): the expression of *ENO1, MIF* and *PRTG* sharply decreases in the healthy branch but remains mostly stable with a downward trend in the DMD branch. In the meantime, the expression of *PDGFRA, ID1* and *PTN* sharply increases in the healthy branch but not in the DMD branch: *PDGFRA* and *ID1* present globally stable RNA counts while the expression of *PTN* decreases; (ii) early switch post-bifurcation (Figure 2d), all 6 gene wave times are less than 6 hours. (iii) strong correlation or anti-correlation with other genes (Figure 2e). In contrast to the wave analysis, the correlation matrix is established between cells from different genotypes, WT versus mutant, independently of their final cell fate. As mutant cells are almost exclusively found in the DMD attractor, we decided to use only WT control cells as they have the potential to end up in the two attractors of interest. The WT correlation matrix indeed shows a clearer typical toggle switch pattern, in contrast with the mutant matrix (Figure 2e). Overall, two pairs of genes - *ENO1*/*MIF* and *ID1*/*PDGFRA* - show a strong positive “intra-” correlation in their expression levels (between *ENO1* and *MIF* and between *ID1* and *PDGFRA*) and a strong negative “inter-” correlation (between the pairs). In other words, cells with a high *ENO1* expression level tend to show high *MIF* expression and low expression of the *ID1*/*PDGFRA* pair of genes.

**Figure 2:**
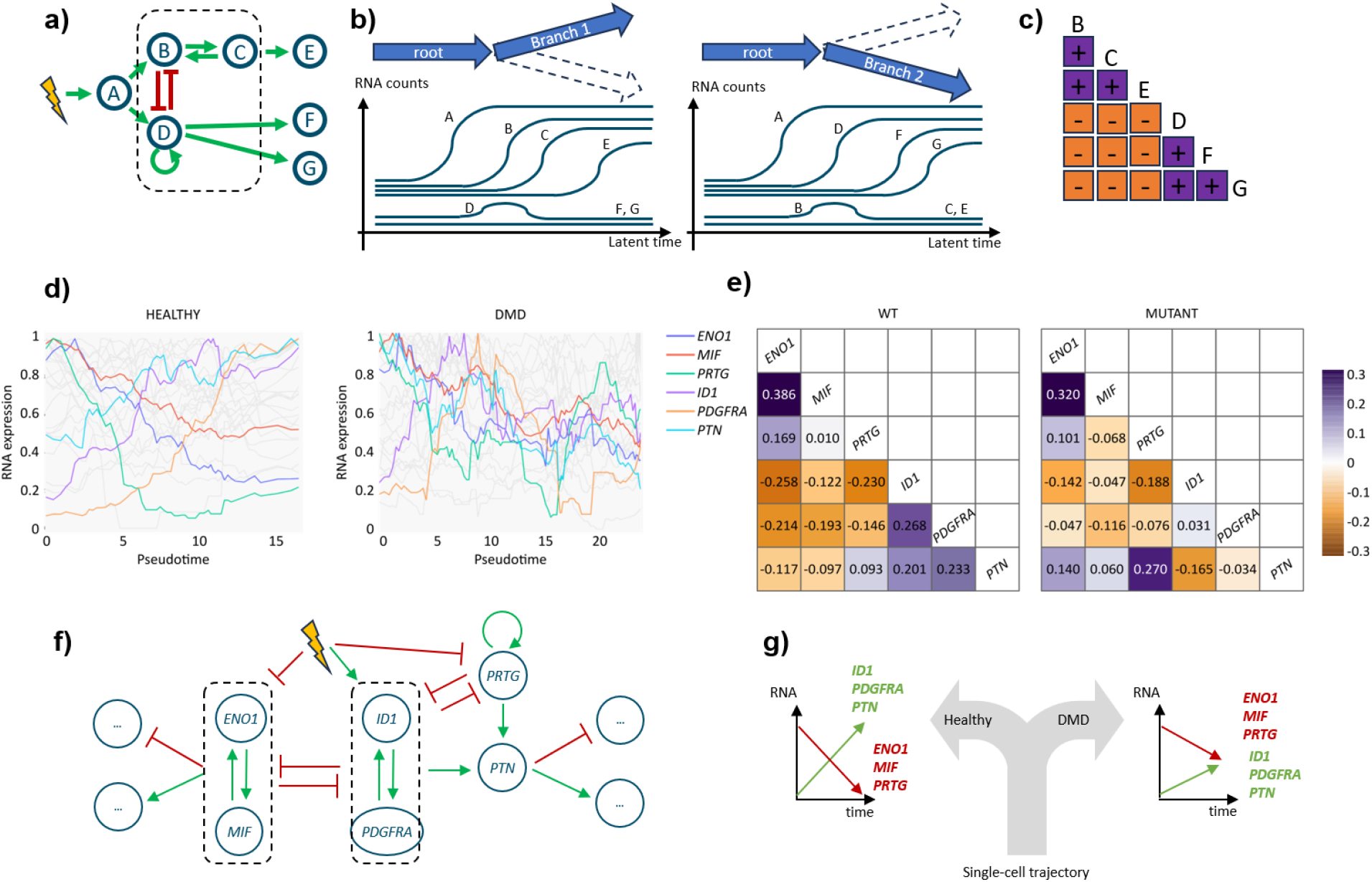
Principle and identification of the core toggle switch GRN. a) Theoretical core toggle switch GRN. Genes B, C and D compose the core toggle switch GRN (dashed rectangle) with 2 kernels in cross-inhibition. The first kernel is composed solely by gene D and its auto-activation, the second kernel is composed by genes B and C with their cross-activation. The toggle switch GRN is indirectly activated by the stimulus through gene A to activate alternatively gene E or genes F and G. b) Theoretical representation of differential waves along both trajectories: Contrary to gene A, toggle switch GRN’s genes and their targets have differential waves along both trajectories. c) Theoretical correlation matrix: Typical toggle switch pattern of a correlation matrix. 2 groups of genes (B, C, E v.s. D, F, G) show positive correlation inside the group and negative correlation between them. d) Experimental waves for each trajectory for the bifurcation trajectory of interest. Experimental conditions WT and mutant are pooled. e) Experimental correlation matrix for each condition (WT, mutant) for the earliest genes with differential waves along trajectories showing a typical toggle switch pattern. Matrix scores correspond to spearman correlation scores. f) Inferred consensus toggle switch network for the WT genotype g) Schematic evolution of toggle switch genes expression during the bifurcation in Healthy and DMD cell fates.

Based on the 6 selected genes we propose a putative GRN with toggle switch features. As such, *ENO1*/*MIF* and *ID1*/*PDGFRA* form two kernels inhibiting each other, which is the basis of a primary toggle switch network (Figure 2f). In addition, *PRTG* creates a second toggle switch with *ID1*/*PDGFRA*, consistent with its negative correlation with this kernel. As *PTN* is not directly involved in kernel topology, we later considered that its participation in cell fate regulation was minimal. The presence of 2 intricate toggle switches wasn’t expected but is compatible with the existence of 4 different attractor states. Therefore, the wave analysis conducted with WASABI led to a consensus GRN (Figure 2f) driving cell bifurcation during the myogenic differentiation of WT cells. This bifurcation is caused by the stochastic activation of a toggle switch, where the expression of *ENO1, MIF* and *PRTG* is reduced while the expression of *ID1, PDGFRA* and *PTN* is increased (Figure 2g). In mutant cells, the activation of the toggle switch is incomplete along the trajectory, leading to an intermediary state, suggesting that the DMD mutation perturbs the toggle-switch gene interactions (Figure 2g).

Importantly, we verified the expression dynamics of toggle switch genes at the RNA and protein levels in bulk. We first confirmed that the protein levels of ENO1, MIF and PRTG follow the trend observed at the RNA level, while ID1 protein count does not (Figure S26). In addition, we performed bulk RNA-Seq and compared the raw expression levels with single-cell RNA-Seq data for the 6 genes of interest. Similar trends were found in both datasets, although bulk data were unable to discriminate WT and mutant cells, suggesting that the toggle switch genes would not have been selected as top targets without data at single-cell resolution.

### Perturbation of the toggle switch partially reverses the DMD cell fate

An important step in systems biology consists in the experimental validation of GRN predicted by in silico analyses. For this purpose, we perturbed the cells experimentally and we generated an independent dataset, as previously published in a previous study involving the WASABI pipeline^18,19^. Here, we aimed to validate the topology of the GRN and its ability to determine cell fate trajectories using the iPSC model. To assess if the toggle switch GRN is at the root of the bifurcation and not a result of upstream – yet undiscovered – regulatory mechanisms, we designed a gene silencing experiment to force the down-regulation of *ENO1, MIF* and *PRTG* in mutant cells using short interfering RNA (siRNA) (Figure 3a). This perturbation is expected to force the switching of the toggle switch genes in mutant cells and consequently to push mutant cells towards the healthy cell fate. Each siRNA was transfected separately in mutant cells at D7 of the differentiation protocol prior to the bifurcation point. Non-transfected and mock-transfected cells were used as controls. We performed a new scRNAseq analysis at D9 and D14 to estimate the impact of siRNA on the mutant transcriptome. The experimental validation protocol is summarized in Figure 3b. To estimate the phenotypic proximity between single-cell transcriptomic data, we first identified 8 marker transcripts that best classify the healthy and DMD states at D9 and D14, independently from the genotype: *COL3A1, GREB1L, SOX2, TMEM132B, LOC105376944, EPHA3, ARHGAP20*, and *NKAIN2*. We then used these markers together with the 6 toggle switch genes for hierarchical clustering of mutant cells transfected with anti-MIF (siMIF), anti-ENO1 (siENO1), anti-PRTG (siPRTG) siRNA and collected at D9 (Figure 3c) and Day 14 (Figure 3d).

**Figure 3:**
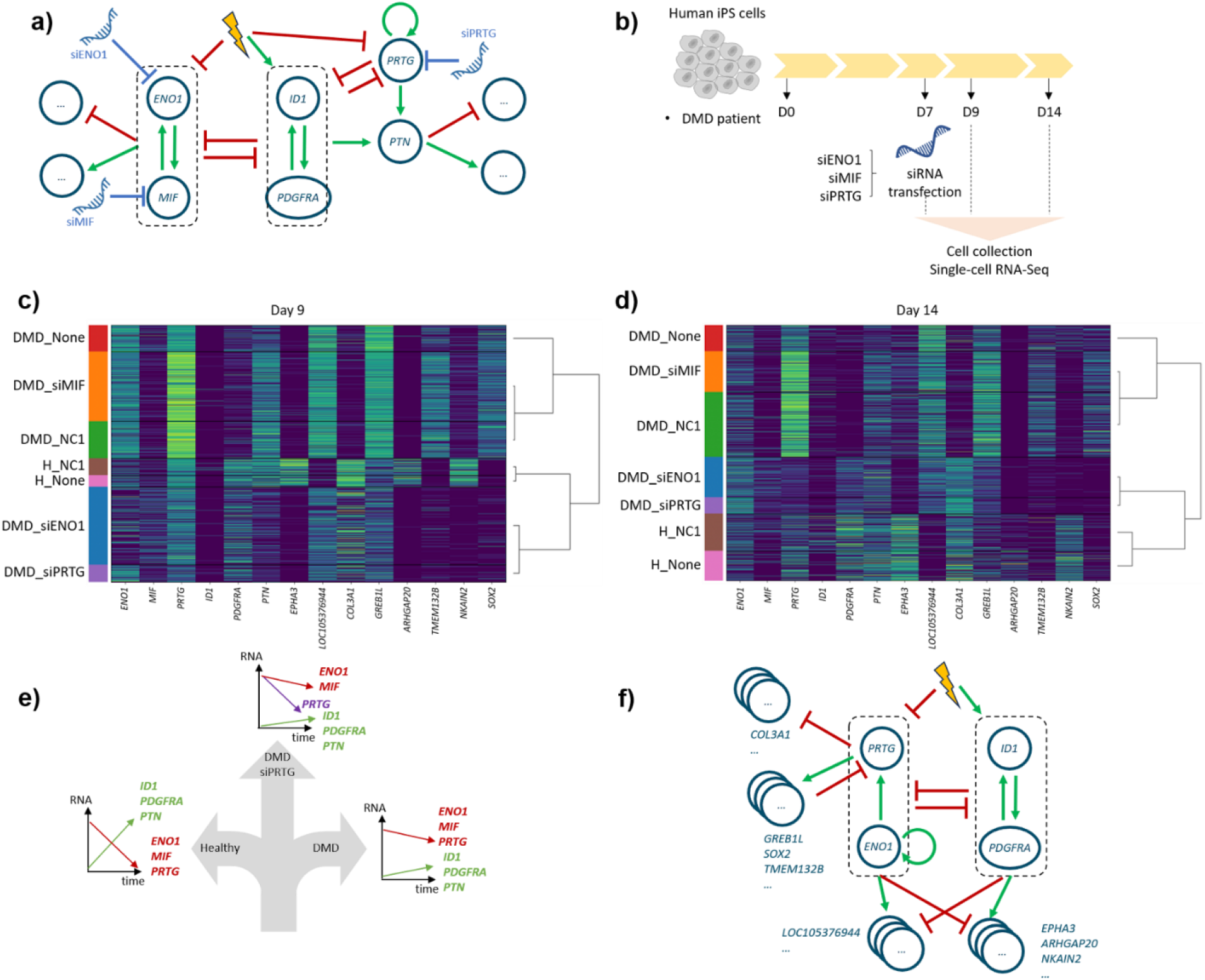
Perturbation of the DMD cell fate and associated GRN by RNA interference. a) RNA interference strategy to force the inhibition of the 3 toggle switch genes active in DMD cells (siENO1, siMIF, siPRTG) and induce a phenotype reversal towards the healthy cell state. b) Experimental protocol of the RNA interference experiment in DMD cells. c, d) Clustering between siRNA treated DMD cells, WT, DMD cells based on scRNAseq data respectively at D9 and D14. Heatmap of RNA expression level for each cell (one cell = one line) grouped by experimental condition (left vertical color bar) for the toggle switch genes and maker genes of the healthy v.s. pathological states (each gene = 1 column). Hierarchical cell clustering based on single cell gene expression. RNA expression heatmap color is normalised regarding maximum expression. e) Schematic evolution of toggle switch gene expression in the context of partial DMD mutation rescue by persistent PRTG inhibition due to siENO1 or siPRTG. f) Updated consensus toggle switch GRN from experimental perturbation results analyses.

As expected, WT cells either non-transfected (H_None), or transfected with a negative control siRNA (H_NC1) cluster together at D9 and D14, suggesting that the transfection protocol does not significantly impact their phenotype. In mutant cells, the effect of siENO1 and siPRTG is visible on *ENO1* and *PRTG* expression, respectively, at D9 and to a lesser extent at D14, suggesting siRNA dilution over time. Unlike our previous data, the expression of *MIF* remains low in all experimental conditions. Interestingly, mutant cells transfected with siENO1 (DMD_ENO1) and siPRTG (DMD_PRTG) cluster together at each time point and then cluster with healthy cells. In contrast, mutant cells transfected with siMIF (DMD_MIF) cluster with mutant cells transfected with the negative control siRNA (DMD_NC1), and both cluster with untransfected mutant cells (DMD_None). Taken together these results suggest that *ENO1* and *PRTG* play a causal role in cell fate determination at the bifurcation point. These results prompted us to update our putative toggle switch GRN topology, in particular the ENO1/MIF kernel that should be replaced by the ENO1/PRTG kernel, which is still consistent with our previous results.

The comparison of gene expression at different times (D9, D14) and perturbations (siENO1, siPRTG) allows us to update the putative gene interactions in the GRN (Figure S27). Focusing on the *ENO1*/*PRTG* kernel, we noticed that *PRTG* expression was reduced with siENO1, but *ENO1* expression was unaffected by siPRTG, suggesting that *ENO1* is upstream of *PRTG*. In addition, *ENO1* expression is partially restored at D14 after siENO1 transfection at D7, suggesting that *ENO1* expression could be restored by a self-activation loop. For the opposite kernel, we observed that the expression of *PDGFRA* – but not that of *ID1* – was increased in DMD_ENO1 and DMD_PRTG cells, which confirms the proposed GRN toggle switch topology where *PDGFRA* is inhibited by the *ENO1*/*PRTG* kernel. Exploring downstream marker genes, we observed a partial rescue of *COL3A1, GREB1L, TMEM123B* and *SOX2* expression in mutant cells transfected with siENO1 and siPRTG, reaching levels comparable to those measured in WT cells. Conversely, the expression of *EPHA3, ARHGAP20* and *NKAIN2* was not significantly affected, while *LOC105376944* expression was only mildly reduced. These results suggest that *COL3A1, GREB1L, TMEM123B* and *SOX2* are only influenced by the *PRTG*/*ENO1* kernel, while *EPHA3, ARHGAP20* and *NKAIN2* should be at least influenced by the opposite kernel and possibly by the *PRTG*/*ENO1* kernel. Strikingly, *PRTG* expression is not restored at D14 with *ENO1* restoration in siENO1 mutant cells, suggesting that *PRTG* could be influenced by a negative feedback induced by its downstream targets. This is a classical GRN pattern in differentiation processes where downstream targets inhibit regulator genes to “erase” the previous cell state^24^. Taken together these perturbation experiments suggest that the persistent inhibition of *PRTG* in mutant cells is sufficient to partially rescue the cell fate and to establish a hybrid state closer to Healthy than DMD (Figure 3e). This iterative experimental validation process enabled us to fine-tune the initial putative toggle switch GRN (Figure 2f) to propose an updated network driving iPSC bifurcation (Figure 3f).

Altogether, our study demonstrates that the DMD cell fate results from an imbalance in a toggle-switch GRN established during disease initiation. The DMD mutation acts as a factor increasing the probability of switching to a pathological state with a specific transcriptomic pattern. In addition, we showed that forced inhibition of toggle-switch genes in mutant cells led to transcriptome remodeling towards that of cells in a healthy state, opening new avenues for early therapeutic interventions.

## Discussion

In the present work, we determined the network of genes regulating cell fate decision in iPSCs submitted to *in vitro* myogenic differentiation, and how it is perturbed by mutations in the *DMD* gene. We described the existence of four attractors among which we distinguished the two distinct cell states - “healthy” and “DMD” - corresponding to the two branches of the single-cell trajectory highlighted in our original study^7^. Importantly, cell fate determination is a stochastic process and WT cells are evenly distributed between the different attractors. In sharp contrast, mutant cells were predominantly found in a single state, suggesting that the DMD mutation unbalances cell fate choice at the bifurcation point.

Of note, the single-cell data generated in the present study have highlighted the existence of two attractors which were not identified in our previous work. This is most likely due to more resolutive data (i.e. higher numbers of cells of each genotype, higher numbers of genes and transcripts detected per cell, and higher number of time points) and to the shorter differentiation protocol focusing more on intermediary differentiation stages.

To understand how the DMD mutation unbalances the cell fate choice probabilities, we adapted the WASABI analysis framework to specifically infer toggle switch genes and their regulatory core networks involved in bifurcation trajectories, based on the estimation of gene expression waves over time. Applied to our dataset, WASABI inferred a GRN embedding a toggle switch-like topology and composed of six core genes. In earlier reports from the literature, we found that some of them are involved in specific functions in skeletal muscle cells. For instance, *ID1* codes for the helix-loop-helix protein ID-1 which sequesters E proteins, regulating the function of basic HLH transcription factors such as MyoD^25^. In turn, ID-1 was shown to inhibit the differentiation of muscle cells^26^. Induction of ID-1 during somitogenesis is a phenomenon already described in fish, allowing to maintain a pool of proliferative progenitors and preventing differentiation into muscle cells^27^. In DMD cells, failure to induce *ID-1* expression at the somite stage might cause progenitors to differentiate prematurely and exit the pool of cells later able to form muscle territories.

Within the GRN, *ID1* forms a kernel with *PDGFRA*, which encodes the platelet-derived growth factor receptor alpha (PDGFRα), a tyrosine kinase receptor with a crucial role in the development of mesodermal tissues^28^. In differentiated muscles, PDGFRα is a marker of stromal cells such as fibroblasts and fibro-adipogenic progenitors, which are key players in DMD pathophysiology^29,30^. During somite development in murine embryos, *Pdgfra* is expressed in the dermatome and sclerotome^31^, and in specific Pax7 progenies which later give rise to dermal fibroblasts^32^. Here, the pathological state is characterized by a defective *PDGFRA* expression in somite progenitors, which could halt their differentiation into dermomyotome derivatives. Overall, the *ID1*/*PDGFRA* kernel seems particularly involved in developmental progression along the skeletal muscle lineage. In DMD cells, the probability to obtain an active kernel is significantly reduced, impairing the efficiency of differentiation into skeletal muscle cells and rather favoring the emergence of aberrant cell types, such as those observed in previous studies by us and others^7,23^.

The second kernel is downregulated when cells bifurcate to the healthy state and involves two genes - *ENO1* and PRTG - activating each other. *ENO1* codes for the alpha-enolase, a glycolytic enzyme expressed in most tissues^33^. In muscle cells, alpha-enolase is also a membrane-bound plasminogen receptor which, in fibrinolytic conditions, allows the accumulation of plasmin and localized protein degradation in the immediate vicinity of myoblasts^34^. This receptor was shown to be required for myotube formation *in vitro* and for muscle regeneration *in vivo*^35^. It can be hypothesized that through its proteolytic activity, alpha-enolase participates in the remodeling of the extracellular matrix necessary for myoblast migration and fusion. On the other side, healthy cells down-regulate the expression of *PRTG* when they reach the bifurcation point. Protogenin (PRTG) belongs to the immunoglobulin superfamily and is well known for its role in axial development. In murine embryos, it was shown to prevent premature neuronal differentiation between E7.5 and E9.5^36^. In the developing somites, a recent study reported that *Prtg*-deficient murine embryos displayed alterations of vertebrae patterning together with aberrant expression of associated Hox genes^37^. Interestingly, delayed expression of posterior *Hox* genes was confirmed *in vitro* with human iPS cells differentiated into presomitic mesoderm.

Importantly, DMD cells treated with siRNAs to force the downregulation of *ENO1* and *PRTG* regain a gene expression profile similar to healthy control cells, at least for specific targets within or directly downstream of the GRN. This confirms that the GRN is modular in DMD cells, and that the pathological cell state does not strictly depend on the mutation, but rather correlates with a specific transcriptomic profile which can be acted upon by exogenous factors. This opens multiple new avenues for future investigations to decipher the gene expression mechanisms at multiple levels, including the epigenetic regulation of promoter activation/inhibition. In particular, the factors sequestered by ID-1 in somite progenitors would require further investigations to better understand the transcription factor dynamics during paraxial mesoderm development and how it is perturbed in a disease context.

DMD is usually diagnosed when children exhibit motor delays at 3 to 4 years of age but disease phenotypes are already present at histological and molecular levels in neonates and fetuses^1,2,38^. Our study uses augmented machine learning for wave analysis coupled with expert-guided feature selection to infer and refine the GRN which determines cell fate during myogenic differentiation of DMD iPSCs, based on the initial transcriptomic profile of each individual cell. The distinguishing feature of our study is based on the experimental validation of the inferred GRN, which not only confirms the biological relevance of WASABI’s wave analysis approach but also sheds light on the mechanisms of DMD initiation at the molecular level.

## Acknowledgements

We thank Dr. Yohan Couté and the EDyP service platform (ISO9001:2015, NFX 50-900:2016 certified, IBISA) for theoretical and technical expertise in proteomics. We thank the GenoA Genomics Core Facility (Nantes Université, SFR Bonamy, UMS Biocore, Biogenouest), which is part of the Institut Français de Bioinformatique (ANR-11INBS-0013) for the access to the computing and storage infrastructure. Additionally, we thank Cyril Seguin, Vincent Lanore ad Cedric Marmonnier for IT support. We also thank Sami Bou-Antoun for his advice and support.

## Author contributions

EM, BR, CF and MC performed the experiments; EGG, QGF, MB, PS, AB and JBD analyzed the data; EGG, QGF, AB and JBD wrote the manuscript and figures; SM, JP, OA, DLM, AB and JBD supervised the study; AB and JBD administered the project; DLM, AB and JBD acquired funding for the project.

## Material & Methods

This study focuses on several experimental datasets. The main single cell RNA seq data (scRNAseq1) was obtained from iPSC lines derived from a healthy donor and a DMD patient. The same differentiation protocol was used to create three new datasets: a bulk RNA sequencing dataset, a proteomics dataset and a single cell RNA sequencing dataset after RNA interference for the validation experiment (scRNAseq2). We will first describe the cell culture procedure which is common to all, then the process for each post culture experimental setup, followed by the data analysis methodology.

### Cell culture

iPSC lines were obtained from a healthy donor and a DMD patient carrying an exon 50 deletion in the *DMD* gene. These lines have been described and published in previous studies^7,39,40^. Informed consent was provided by the participants as required by the Institutional Review Board of the University of Washington. The experiments described in this manuscript are part of Category 1A from the “Guidelines for Stem Cell Research and Clinical Translation” of the International Society for Stem Cell Research (iSSCR, https://www.isscr.org/guidelines). Briefly, urine samples were collected from the participants and urine stem cells were isolated for iPSC generation by Xuan Guan at the University of Washington (Seattle, USA). The resulting iPSC lines were then shared with the TaRGeT laboratory at Nantes Université (Nantes, France) after signing a Material Transfer Agreement (MTA).

iPSC lines and their derivatives were maintained in standard culture conditions at 37°C and 5% CO2. Master Cell Banks (MCB) were generated from parent iPSC vials at specific passages (Healthy: P34; DMD: P25) and used for Working Cell Bank (WCB) preparation after two passages. For this purpose, iPSCs were rapidly thawed at 37°C and plated as clusters on matrigel-coated 6-well plates (CORNING, NY, USA, dilution 1:60 in DMEM/F12) after spinning 3 min at 300 g. They were first cultured overnight in mTeSR Plus medium (STEMCELL Technologies, Vancouver, Canada) supplemented with 10 µM ROCK inhibitor (EUROMEDEX, Strasbourg, France) to maximize iPSC attachment and survival. The day after, the culture medium was replaced by fresh mTeSR Plus without ROCK inhibitor. Medium was changed every other day after visual inspection and manual removal of abnormally looking clusters. When iPSC clusters reached 70 to 80% confluence, they were gently passaged using Versene (Thermo Fisher Scientific, Waltham, USA) and mechanical dissociation with a cell scraper. After two passages, iPSC clusters were frozen in Cryostor freezing media (STEMCELL Tehcnologies) and WCB vials were long-term stored in liquid nitrogen.

iPSCs were differentiated into paraxial mesoderm derivatives using a previously published protocol (Chal 2016). Briefly, iPSCs were seeded as single-cells in 1:60 matrigel-coated 12-well plates at a density of 30,000 cells/cm^2^ using mTeSR Plus supplemented in ROCK inhibitor (10 µM) as culture medium. Two to three days later, differentiation was initiated and continued for 15 days.

Five successive culture media were used:

- M1 (3 days, daily change): DMEM/F12 (Thermo Fisher Scientific), non-essential amino acids (NEAA, Thermo Fisher Scientific) 1X, Insulin transferrin selenium (ITS, Thermo Fisher Scientific)) 1X, CHIR99021 3 µM (STEMCELL Technologies), LDN 0.5 µM (Miltenyi Biotec, Bergisch Gladbach, Germany).
- M2 (3 days, daily change): M1 supplemented with basic Fibroblast Growth Factor (bFGF, R&D Systems, Bio-Techne, Minneapolis, USA) at 20 µG/ml
- M3 (2 days, daily change): DMEM/F12, non-essential amino acids (NEAA) 1X, 15% Knockout serum replacement (KSR, Thermo Fisher Scientific), LDN 0.5 mM, bFGF at 20 mg/mL, Hepatocyte Growth Factor (HGF, R&D Systems) at 10 mg/mL, Insulin-like Growth Factor (IGF, R&D Systems) 1 at 2 ng/mL
- M4 (4 days, daily change): DMEM/F12, non-essential amino acids (NEAA) 1X, IGF1 at 2 ng/ml
- M5 (3 days, change every other day): M4 + HGF at 10 ng/ml.

### scRNAseq1

Cells were filtered at 70µM and washed (500g, 5min, 4°C) in PBS then counted and checked for viability. They were resuspended in 100 uL of staining buffer (1xPBS with 2%BSA, 0,01% Tween20, 0,22µM filtered) and incubated with 10 µL human FcR blocking reagent (Miltenyi Biotec 130-059-901) for 10 min on ice. 1 µL of anti-human Totalseq-B Hashtag oligonucleotides antibodies (HTOs) (Biolegend, San Diego, USA) were assigned to each sample and incubated for 30 min on ice. Tagged cells were then washed three times in 1mL of staining buffer (500g, 5min, 4°C). Cells were counted and checked for viability, then pooled in equal proportions per sample. The viability of the pools varied between 71-98%. A total of 20000 cells per pool were loaded on the Chromium Next Controller using the Chromium Next GEM Single Cell 3’ GEM, Library & Gel Bead Kit v3.1 (PN-1000121, 10x genomics, Pleasanton, USA). We followed the supplier’s user guide CG000206 Rev D and generated 3’GEX and Cell Surface (CS) libraries using 3’ feature barcode kit (PN-1000079) during pre-amplification step. For PCR index, we used Single Index Kit T Set A, 96 rxns PN-1000213 for 3’GEX CS. All libraries were checked on a Bioanalyzer with a High Sensitivity chip (Caliper Perkin Elmer, Waltham, USA). As expected, 3’ GEX libraries peaked around 350pb and CS at 220pb. Libraries were then sequenced on a Illumina NOVAseq 6000 (Illumina, San Diego, USA), on a 100-cycle S1 v1.5 flow cell kit (Illumina) with the R1-29pb/i5-0pb/i7-8pb /R2-93pb running program at the Genomic Atlantic (GenoA) platform (IRS-UN, CHU Nantes, France).

### scRNAseq2

Commercial siRNA preparations (Integrated DNA Technologies, Coralville, USA) were used and transfected in iPSCs at Day 7 of myogenic differentiation using RNAiMAX lipofectamine (Thermo Fisher Scientific). Differentiation media supplemented with ROCK inhibitor at 10 µM was changed at least 2 hours before transfection. Cells were dissociated with Trypsin-EDTA (Thermo Fisher Scientific) supplemented with collagenase IV at 50 U/ml (Thermo Fisher Scientific) after incubation for 10 min at 37°C. The cell suspension was filtered through a 70 µm strainer and Trypsin was inactivated by addition of DMEM/F12 supplemented with 10% fetal bovine serum. After spinning at 1000 g for 5 min, aliquoting and trypan blue addition, living cells were counted with an automated cell counter and divided in separate tubes with 2.0E+6 cells per tube. The siRNA were diluted at 72 µM in OptiMEM medium (Thermo Fisher Scientific) and mixed with lipofectamine RNAiMAX diluted at 1:10 in OptiMEM. The tubes were incubated 45 min at RT min and then used to resuspend 2.0E+6 cells. The transfection mix was incubated at RT for 10 min and diluted with 1.9 ml of differentiation media 3 + ROCK inhibitor 10 µM before seeding on a matrigel-coated 6-well plate. The medium was changed the day after transfection and differentiation was pursued normally. Samples collected after siRNA transfection were dissociated with Trypsin-EDTA supplemented with collagenase IV at 50 U/ml after incubation for 10 min at 37°C. Preparation for split-pool barcoding first involved a Fixation-Permeabilization step using the Evercode™ Cell Fixation Kit (v2, Parse Biosciences, Seattle, USA). Fixed cells were kept at −80°C until library preparation. Samples were thawed and counted with an automated cell counter (Countess II, Thermo Fisher Scientific). Approximately 12,000 cells of each sample were used as inputs and dispatched into the first reaction plate of the Evercode™ Whole Transcriptome Kit (Parse Biosciences). The kit was then used according to the manufacturer’s instructions. The library was sequenced on a Illumina NOVAseq 6000 (Illumina), on a 200-cycle Sp flow cell kit (Illumina) with the R1-108pb/i5-0pb/i7-6pb /R2-86pb running program at the Genomic Atlantic (GenoA) platform (IRS-UN, CHU Nantes).

### Bulk RNA seq

At Days 5, 7 and 9 of muscle differentiation, iPSCs were treated with actinomycin D (Sigma Aldrich, Saint-Louis, USA) at a final concentration of 10 µg/m. Cells were then collected at 1, 2, 3 and 4 hours after treatment and 1.0E+4 to 1.0E+6 cells were pelleted and snap frozen. RNA extraction and bulk sequencing was performed by Genewiz (South Plainfield, USA). After quality control with fastQC, the bulk RNA seq dataset was aligned using kallisto^41^. From this dataset, with inhibition of activation we get a count of transcript for each gene at each timepoint. Using the time of the inhibition only we get the dynamic evolution of RNA in DMD and healthy states. This study did not use the additional data points.

### Proteomics

Cells were collected at 8 successive time points: Days 5, 6, 7, 8, 9, 10, 12 and 14. After a wash with DPBS-1X, differentiating cells were incubated with Trypsin-EDTA 0.25 % for 7 min at 37°C. After Trypsin neutralization with DMEM-F12 10 % FBS, cells were centrifuged at 300g for 5 min and counted with a hemocytometer. Samples containing 2.0E+6 cells were transferred in cryotubes and frozen at −80°C. They were shipped to the EDyP Service platform (Grenoble, France) for nano-Liquid Chromatography and Mass Spectrometry (nano-LC-MS/MS) after sample preparation with the iST-NHS kit (PreOmics, Planegg, Germany) and TMTpro 16plex Label Reagent Set (Thermo Fisher Scientific).

### scRNAseq data analysis

The quality of the scRNAseq reads was checked using the FASTQC software (http://www.bioinformatics.bbsrc.ac.uk/projects/fastqc). All indicators of quality were within their expected range. Cross-species contamination was checked using the Fastq Screen software^42^. Reads were aligned on the human genome GRCh38 using CellRanger v7.1.0 (10X Genomics) to obtain bam files. Velocyto^43^ was then used to align the reads, directly from these bam files to the human transcriptome. After alignment, cells with a low number of UMI and a low number of genes counts were removed using an adjusted threshold for each sequence batch based on the UMI/cell and the genes/cell distributions, as is standardly practiced^44^. No evident batch effect was identified across the two sequencing runs (Figure S1, S2). Other metrics, such as percentage of mitochondrial and ribosomic RNA were checked, but not used to filter out cells. Data were normalized, genes with less than 10 total counts in the samples were filtered out. All scRNAseq data were processed with scanpy^45^. After normalization, there are 10,965 cells in the dataset scRNAseq1, details are given in the supplementary material (Table S1).

### Trajectory analysis

For trajectory analysis, the dataset including all cells and genes was projected in two dimensions using the UMAP algorithm^21^. Figure 1d shows cells ordered following their time point with a progressive expansion through time. Two methods are used to compute the cell velocity: the stochastic model (Figure S3) and the dynamical model (Figure S4) from scvelo^46^, the latter one was used for the rest of the analysis. Phenotyping of the cells was necessary to perform feature selection. Based on a list of genes (Table S2) marking four different stages of muscle differentiation (pluripotency -> mesoderm -> somite -> muscle), the AUC score^22^ was computed for each stage to annotate the cells on the UMAP (Figure S5). The first feature selection focuses on cells from day 9 to day 14 (Figure S9) followed by the removal of pluripotent cells identified by cell cycle scoring^47^ (Figure S12-S14). On this subset we perform again a UMAP dimension reduction and velocity computation. Using the latest as well as the score associated with muscle differentiation, we identified 4 cell subsets that characterize the target trajectory (Figure S14). To study the trajectory, we reduced the dataset to genes that were differentially expressed between these cell subsets (t-test with BH correction, log2 fold change > 2 and a qval < 0.01), keeping 1,295 genes. After UMAP and velocity was performed on this subset (Figure S15), the trajectory identified on the new projection (Figure S16, 17) we computed the latent time along the trajectory using scvelo^46^ (Figure S20).

### GRN inference

On the trajectory of interest with associated latent-time, we computed RNA gene waves according to the original method described in the WASABI algorithm^17^. The 698 waves obtained were then filtered by slope (between −20 and 10 counts / hour), relative variation (>0.15) and stabilization time (>0.25 hour). This resulted in a subset of 140 genes with the most significant waves. We hypothesized that the bifurcation trajectory is driven by a GRN with a toggle switch topology, which was inferred based on the analysis of waves using the WASABI framework. Figure 2a represents a typical toggle switch network composed of 3 genes: B, C and D. We call “kernel” the subnetwork formed by genes B and C. In the example depicted in Figure 2a, genes B and C create a kernel with an auto-activation loop, while gene D constitutes the second kernel and has its own auto-activation loop. During a differentiation process, genes are activated/repressed due to specific stimuli and gene interactions. If a toggle switch pattern is present in the network under the influence of the stimulus, it creates a bifurcation in the phenotypic landscape. Figure 2b represents a theoretical bifurcation trajectory driven by the gene regulatory network of Figure 2a, and the associated waves of gene expression. Waves are the central concept of the WASABI inference method. They correspond to the transient upregulation or downregulation of genes submitted to an upstream regulation. If we follow the gene expression along a latent time for cells starting from the “root” state and undergoing the bifurcation, we could observe distinct wave patterns in trajectory 1 and trajectory 2. In trajectory 1 represented in Figure 2b, genes B, C, E are activated and genes D, F, G stay inactivated, and vice-versa in trajectory 2. Thus, genes involved in the toggle switch, or under the influence of the toggle switch, have differential patterns in both trajectories. Conversely, Gene A has the same activation pattern in both trajectories because it is not influenced by the toggle switch. Another characteristic to identify the toggle switch genes is the typical correlation matrix pattern shown in Figure 2c. Genes involved in a toggle switch or under its influence create 2 groups, called kernels, with intra-positive-correlations (positive correlation between the genes of the same kernel), and inter-negative-correlations (negative correlation between the genes of different kernels). Altogether, the genes involved in the toggle switch must fulfill 3 conditions: 1) to have differential wave patterns between trajectories 2) to be the earliest waves among differential waves, 3) to present a typical toggle switch correlation matrix pattern.

### Marker genes

Marker genes for the DMD and healthy states are inferred using the scRNAseq2 dataset. A subset of the data with cells at day 9, mutant and WT, either non-transfected or transfected with the negative control siRNA NC1. Random forest algorithm was used to learn cell classification in either healthy or DMD based on all genes in the dataset, using 80% of cells as tests and 20% as control. The top 8 genes with a significance > 0.1 were used as markers of the healthy and dmd states. They are plotted in Figure 3c and 3d alongside genes found in the trajectory analysis as core regulatory genes.

## Data availability

The datasets produced in this study will be made available in the following database:

- RNA-Seq data: Gene Expression Omnibus

