## Supplemental Material for "Inference and validation of a dynamic gene regulatory network influencing cell fate during Duchenne muscular dystrophy initiation"

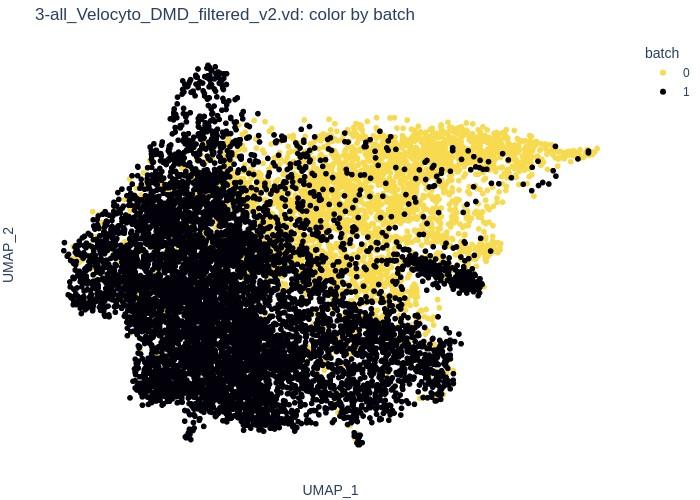

Figure S1: UMAP on the whole scRNAseq1 dataset colored by batch (0 and 1).

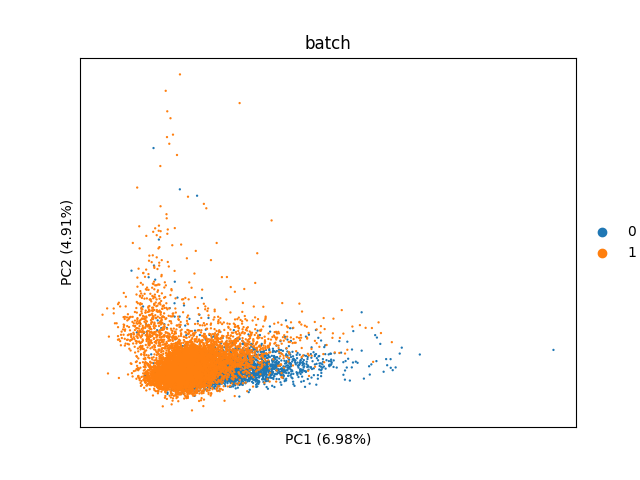

Figure S2: PCA representation of scRNAseq1 data coloured by batch. With a low value of PC1 and a large overlap of cells, this shows that there is no batch effect. The small shift observed is due to the time effect that is confounded with the batch effect.

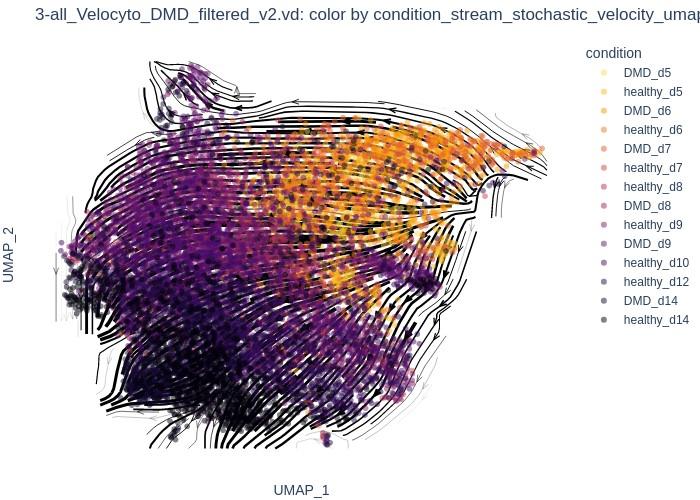

Figure S3: UMAP representation of the whole scRNAseq1 dataset with stochastic RNA velocity and colored by condition (DMD or healthy, from day 5 to day 14).

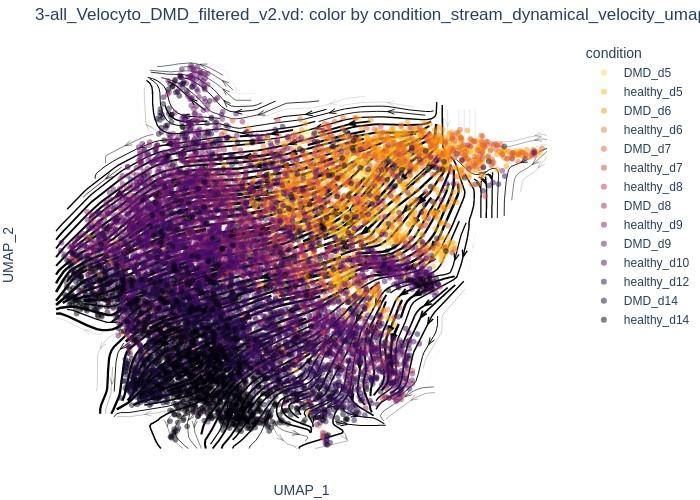

Figure S4: UMAP on the whole scRNAseq1 dataset with dynamical RNA velocity and colored by condition (DMD or healthy, from day 5 to day 14).

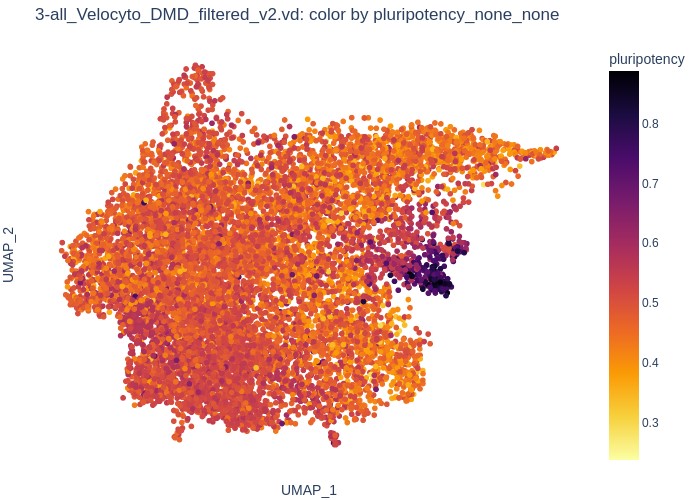

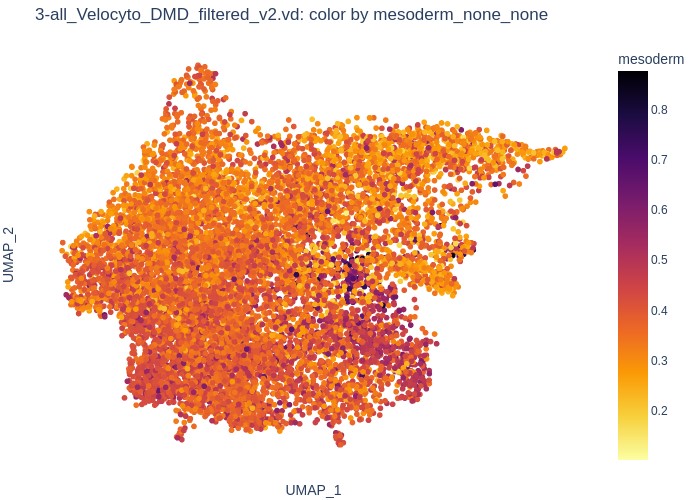

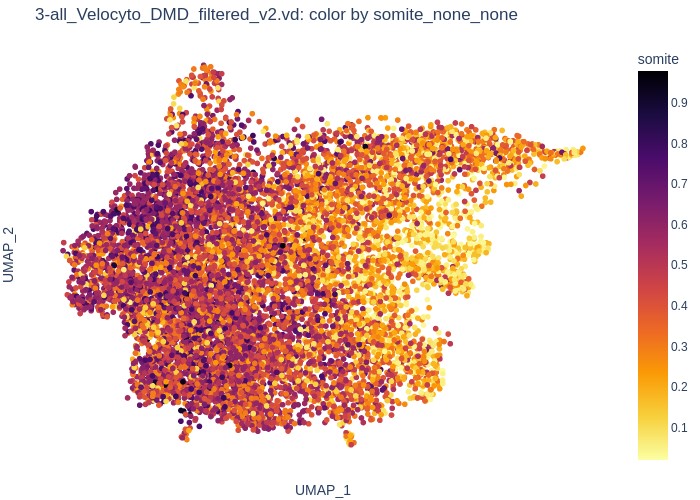

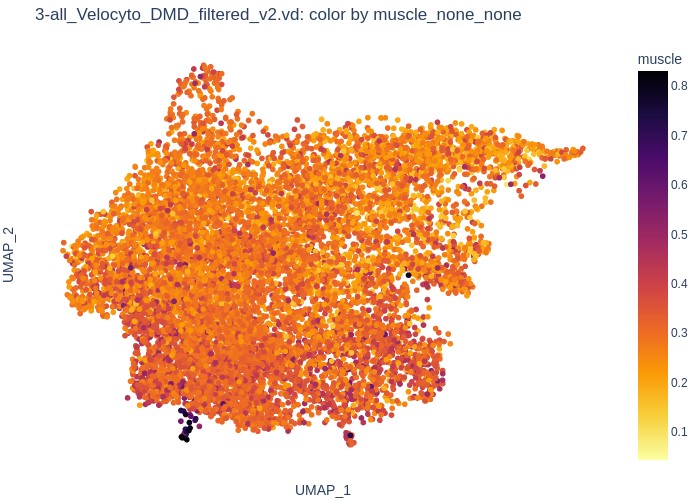

Figure S5: UMAP representation of the scRNAseq1 dataset colored by AUC score for successive differentiation stages (pluripotency, mesoderm, somite and muscle). For more information on AUC scores, see Aibar, S. et al. SCENIC: single-cell regulatory network inference and clustering. Nat Methods 14, 1083–1086 (2017). Reference 22 of the main manuscript.

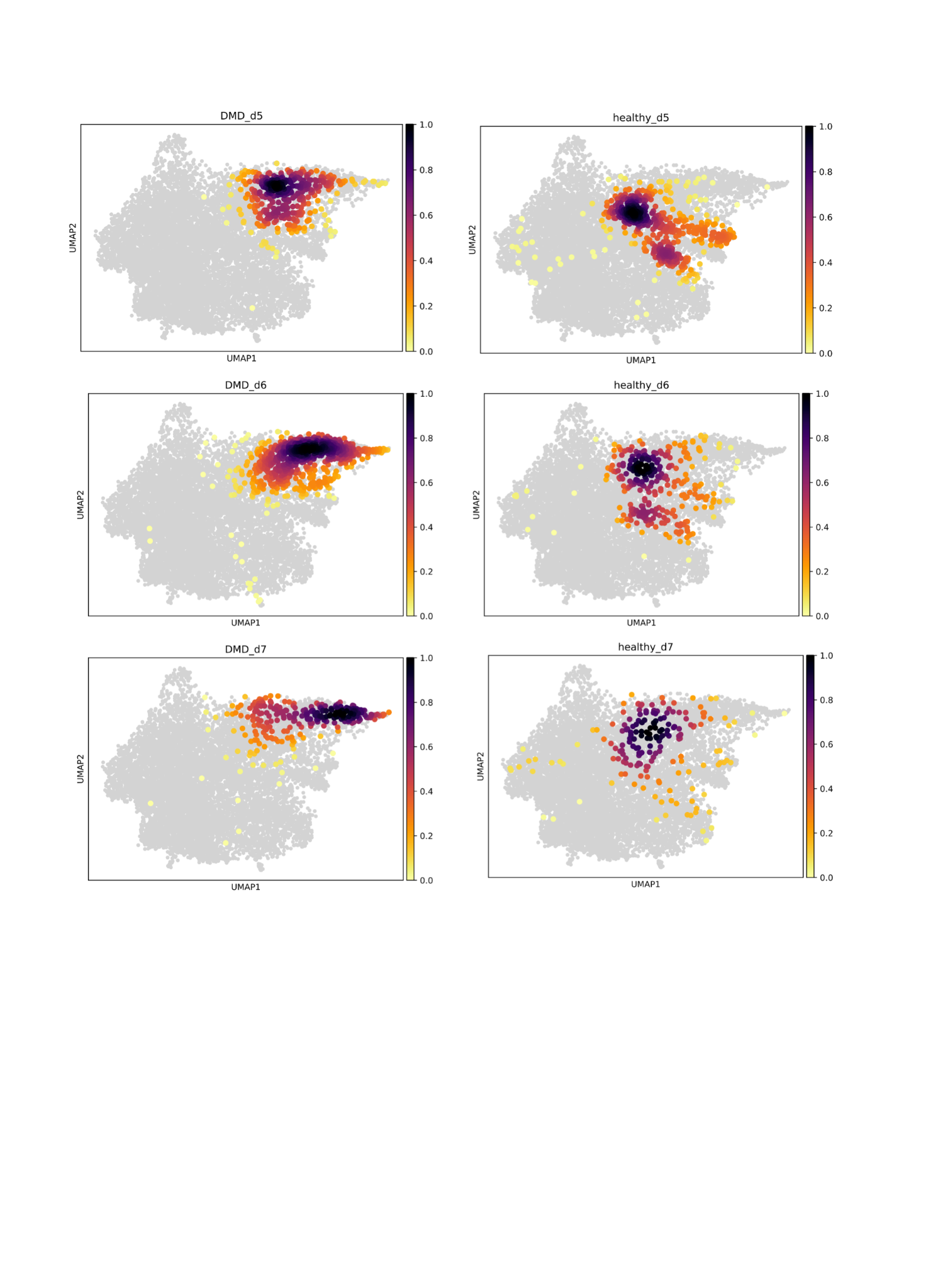

Figure S6: UMAP representation of the scRNAseq1 dataset colored by cell density for each experimental condition (DMD and healthy), between day 5 (d5) and day 7 (d7).

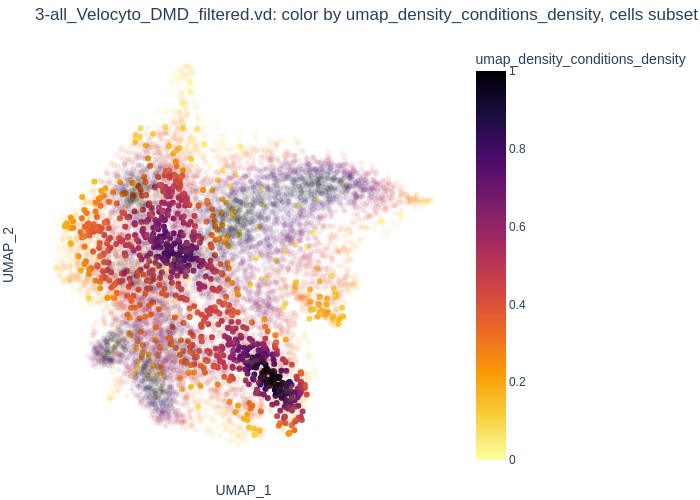

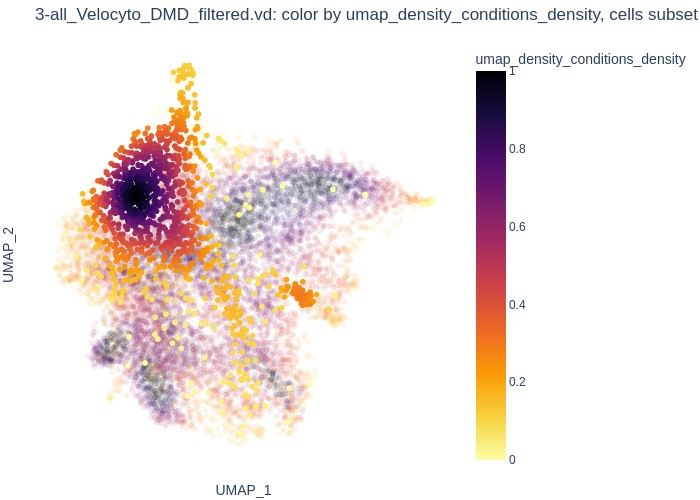

(a) healthy day 9	(b) DMD day 9

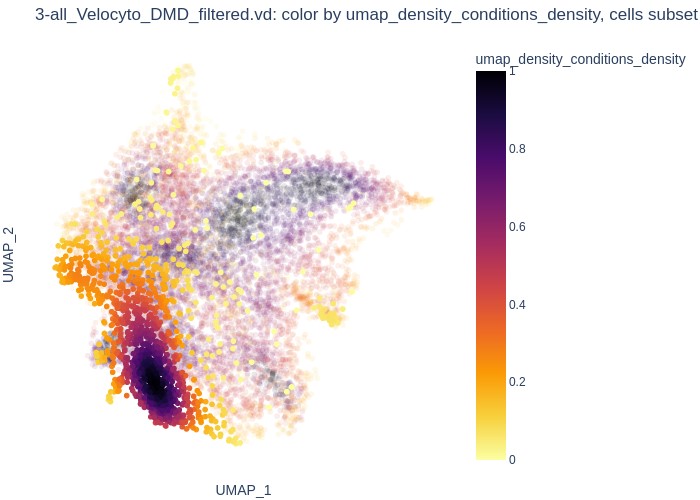

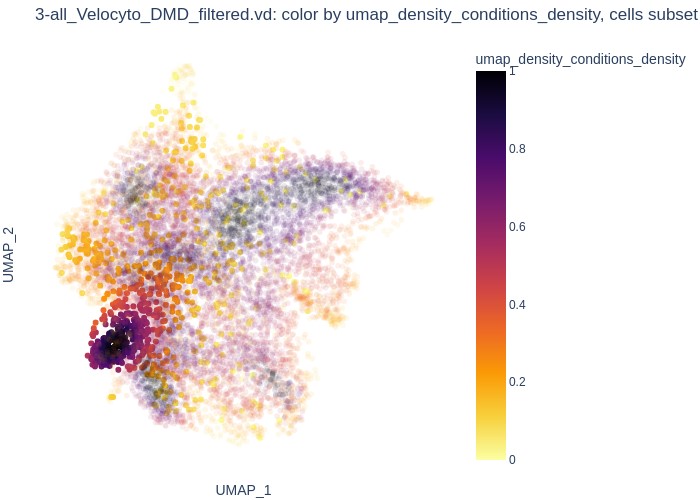

(c) healthy day 14	(d) DMD day 14

Figure S7: UMAP representation of the scRNAseq1 dataset colored by cell density for each experimental condition (DMD and healthy) at day 9 (d9) and day 14 (d14).

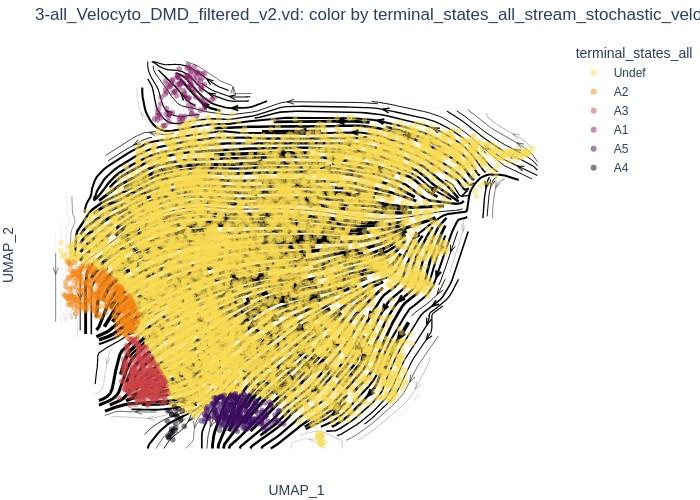

Figure S8: UMAP representation of the whole scRNAseq1 dataset with dynamic RNA velocity data colored by terminal states.

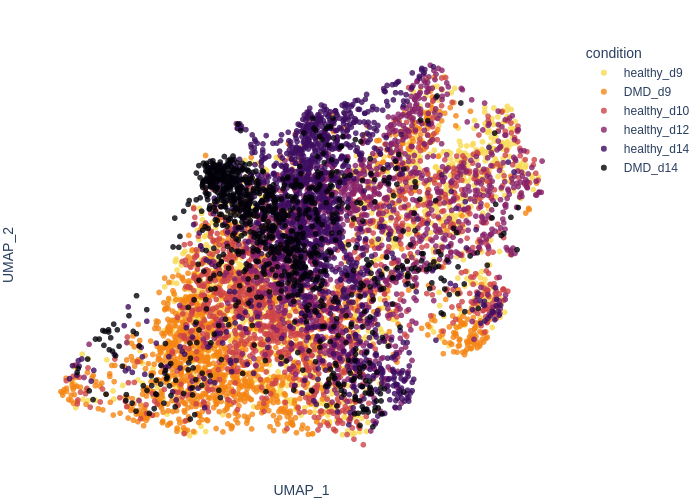

Figure S9: UMAP representation of the scRNAseq1 dataset after subset of the samples collected between day 9 (d9) and day 14 (d14) for the healthy and DMD cells.

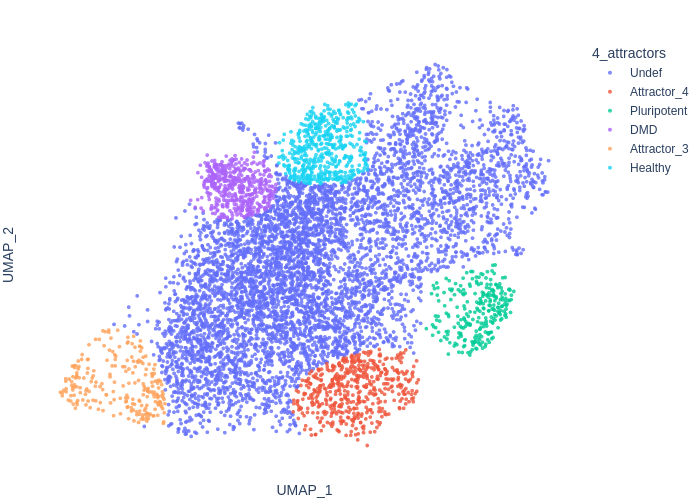

Figure S10: UMAP representation of the scRNAseq1 dataset with a subset of the samples collected between day 9 (d9) and day 14 (d14) and colored by cell states.

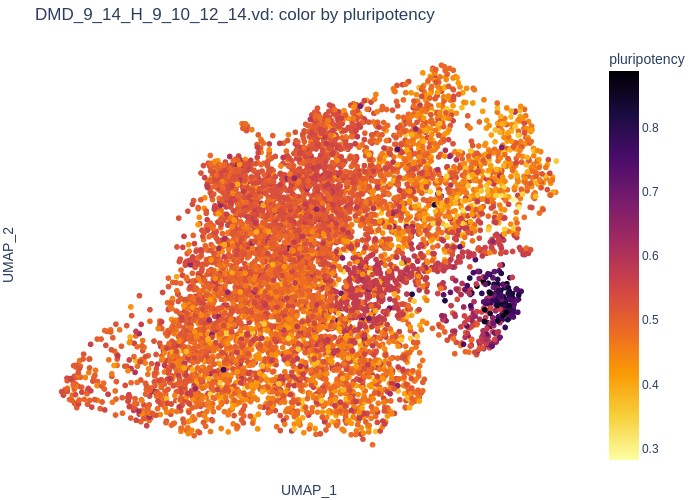

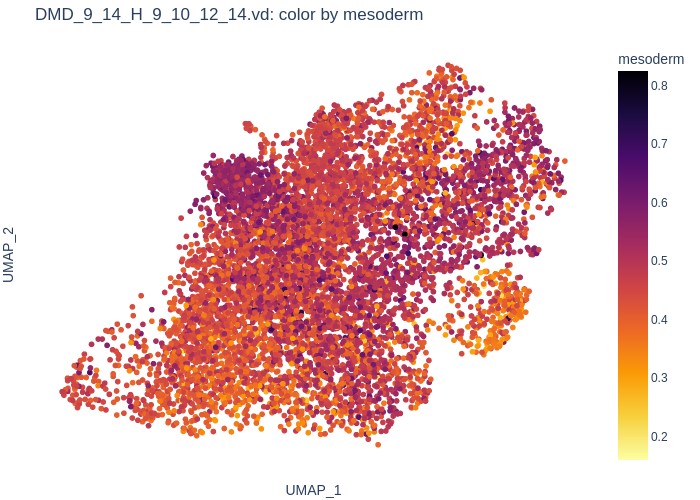

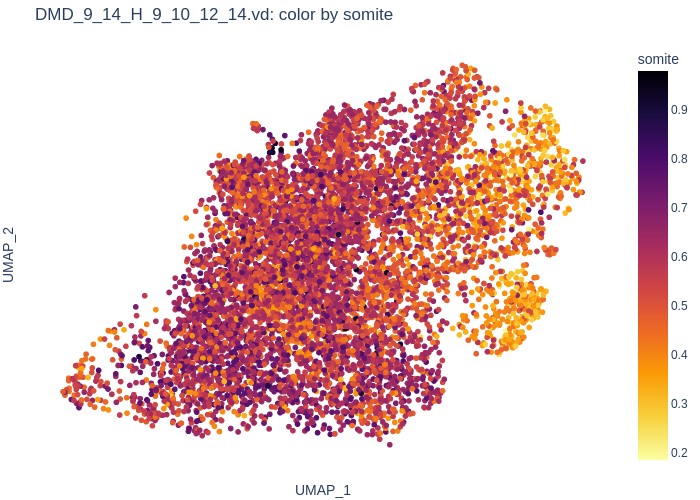

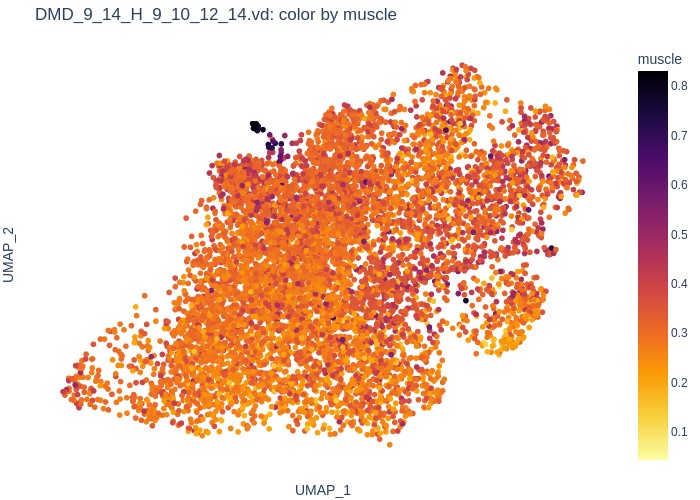

Figure S11: UMAP representation of the scRNAseq1 dataset, subset of samples collected between day 9 and day 14, and colored by AUC score for successive differentiation stages (pluripotency, mesoderm, somite and muscle). For more information on AUC scores, see Aibar, S. *et al.* SCENIC: single-cell regulatory network inference and clustering. *Nat Methods* **14**, 1083–1086 (2017). Reference 22 of the main manuscript.

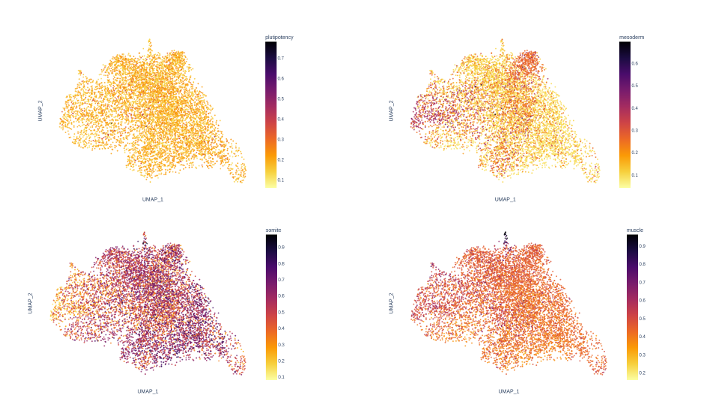

Figure S12: UMAP representation of the scRNAseq1 dataset, subset day 9 to 14, with pluripotent cells removed and colored by AUC score for successive differentiation stages (pluripotency, mesoderm, somite and muscle). For more information on AUC scores, see Aibar, S. *et al.* SCENIC: single-cell regulatory network inference and clustering. *Nat Methods* **14**, 1083–1086 (2017). Reference 22 of the main manuscript.

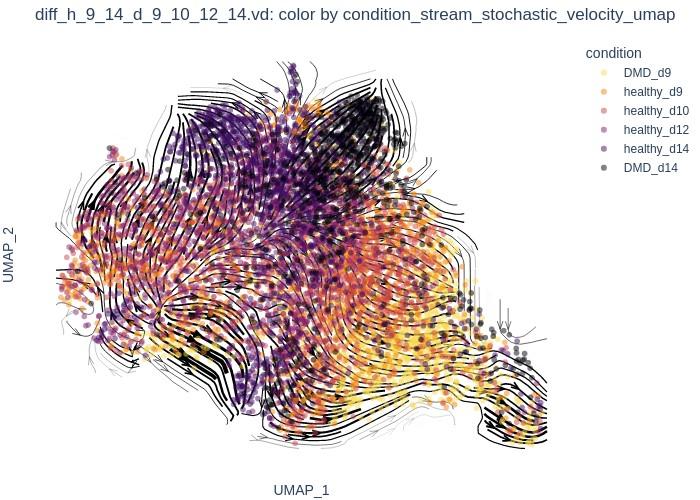

Figure S13: UMAP representation of the scRNAseq1 dataset with subset of samples collected between day 9 and day 14, removal of pluripotent cells, and colored by condition. Stochastic RNA velocity represented with black arrows.

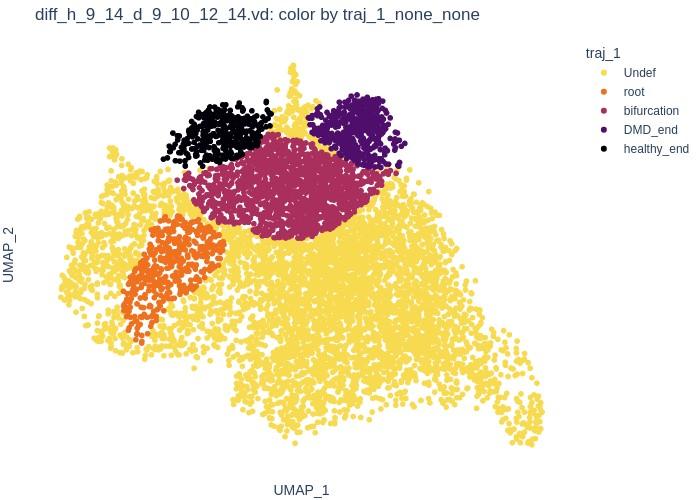

Figure S14: UMAP representation of the scRNAseq1 dataset with subset of samples collected between day 9 and day 14, removal of pluripotent cells, and colored by the set_of_trajectory: ’traj_1’ cell attribute.

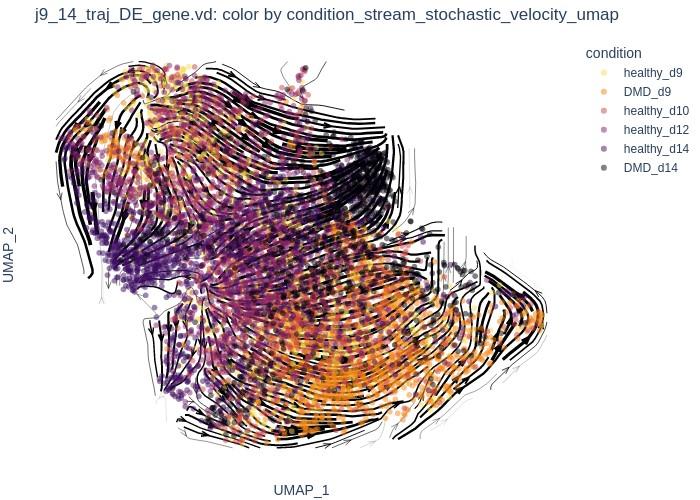

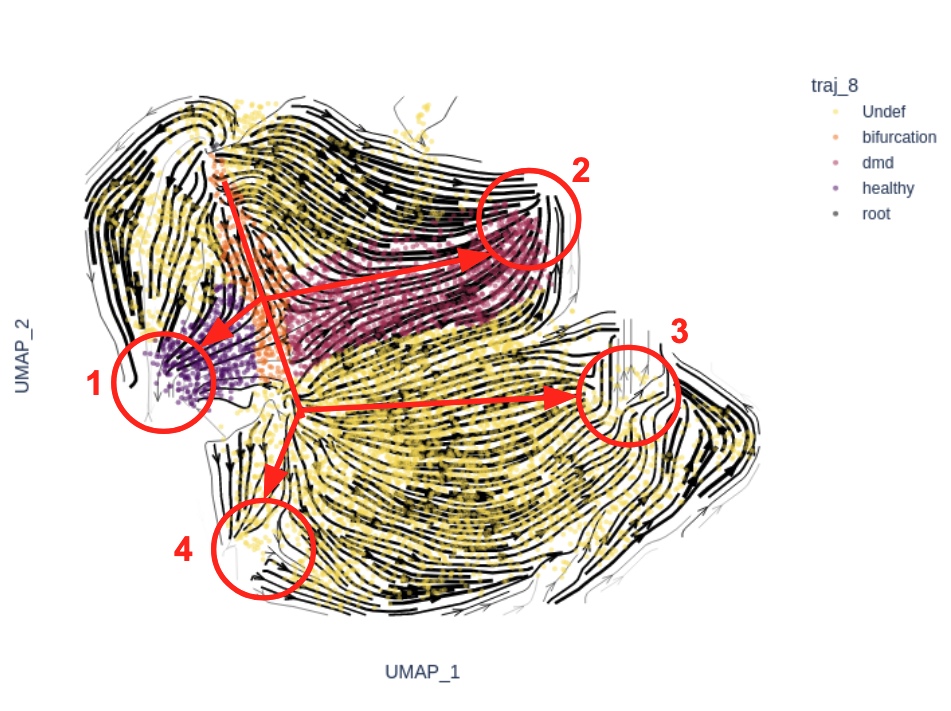
Figure S15: UMAP representation of the scRNAseq1 dataset with subset of samples collected between day 9 and day 14, removal of pluripotent cells, and restriction to differentially expressed genes between clusters. Cells are colored by conditions (Undef, bifurcation, dmd, healthy and root). Stochastic RNA velocity is represented with black arrows.

Figure S16: Bifurcation trajectory highlighting with the 4 final attractors. Attractors 1 and 2 correspond to the two final cell states of interest. Attractor 3 corresponds to “neural development” while attractor 4 is composed of cells engaged in the G2M phase of the cell cycle.

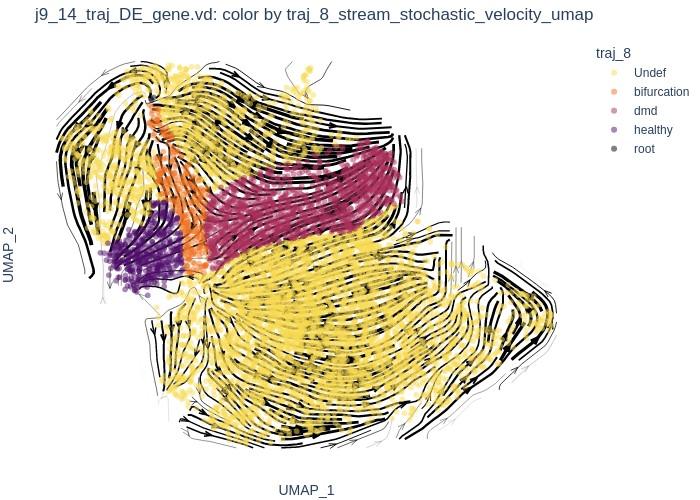

Figure S17: UMAP representation of the scRNASeq1 dataset after subset of samples collected between day 9 and day 14, removal of pluripotent cells, restriction to differentially expressed genes between clusters and definition of the genetic trajectory of interest, composed of one bifurcation state and two final attractors: healthy and DMD.

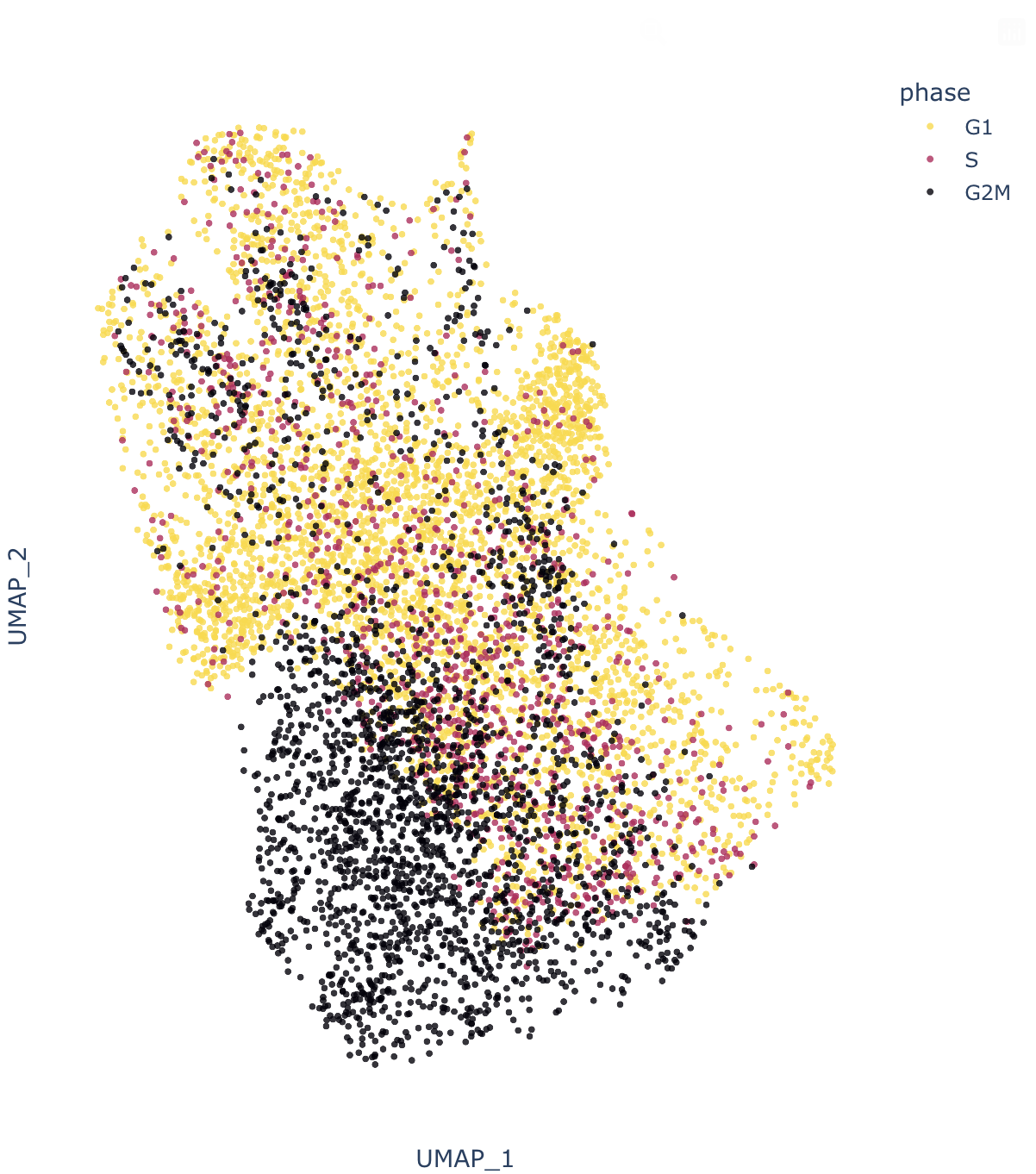

Figure S18: UMAP representation of the same dataset as in Figure S17 and showing the cell cycle phase as inferred using scanpy. This shows an enrichment in G2M cells in the bottom left attractor (attractor 4), suggesting that these cells are engaged in mitosis.

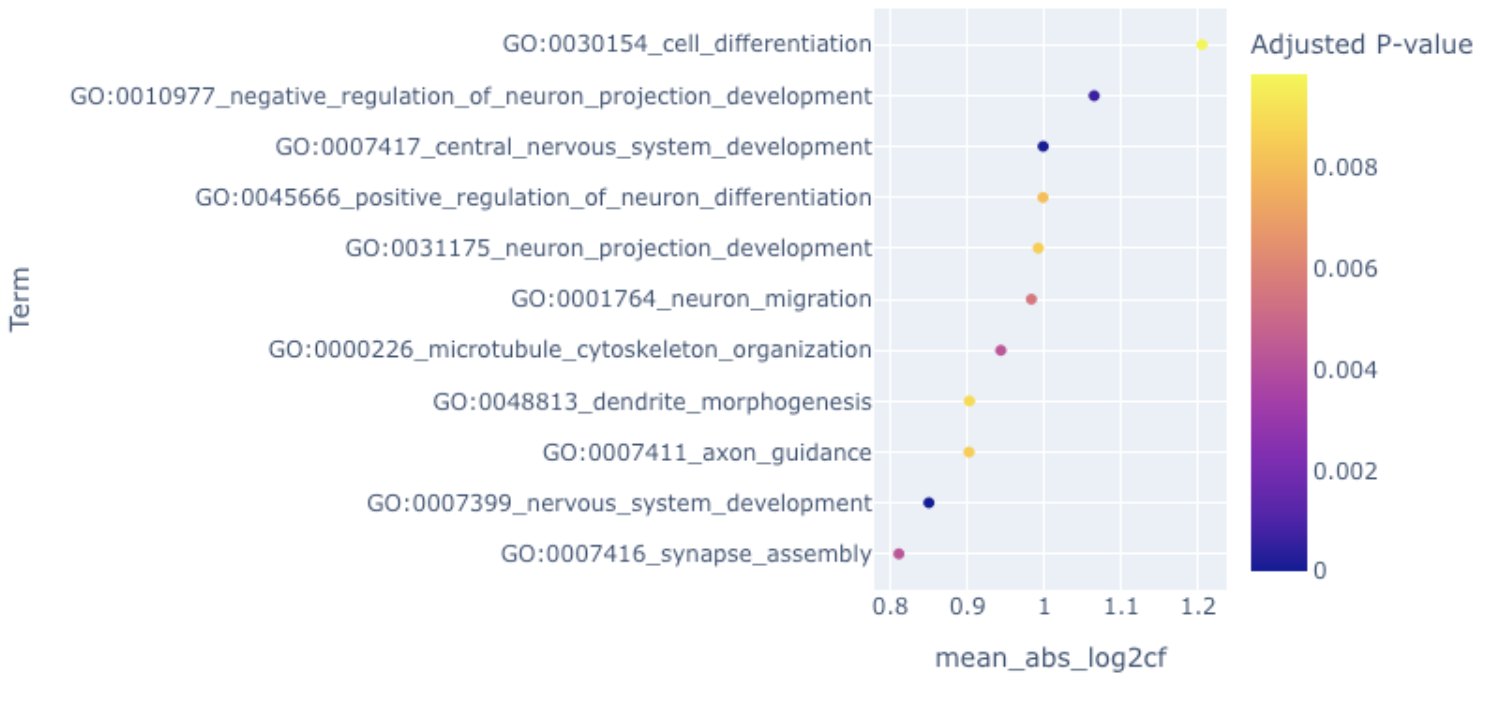
Figure S19: Gene set enrichment analysis of genes that are differentially expressed in the attractor “neuron development” with a q-value < 0.01 and log fold change > 2 using GO terminology. This shows an enrichment in genes involved in neural development processes.

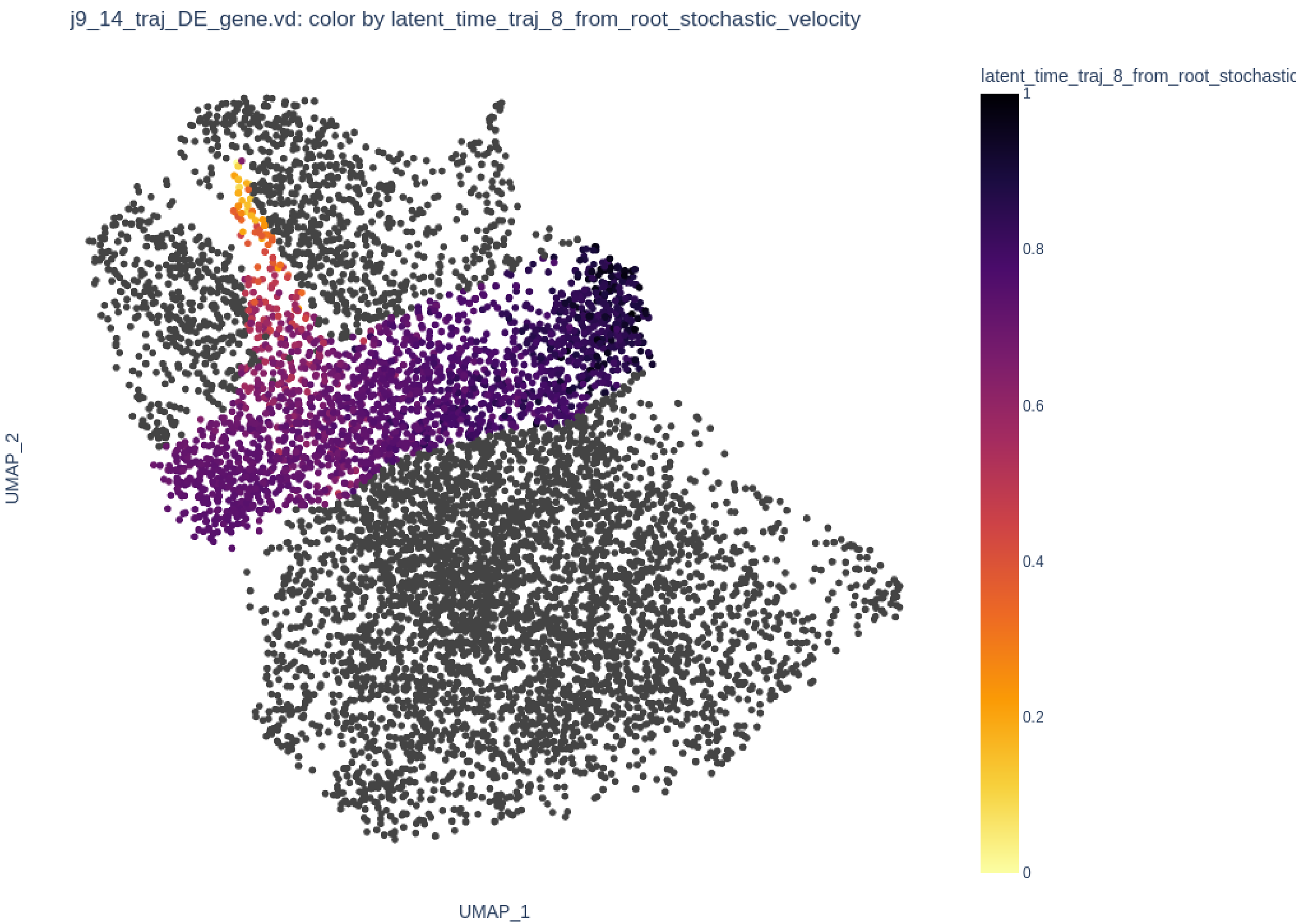

Figure S20: UMAP representation of the same dataset as in Figure S17 and showing the latent time evolution along the trajectory of interest.

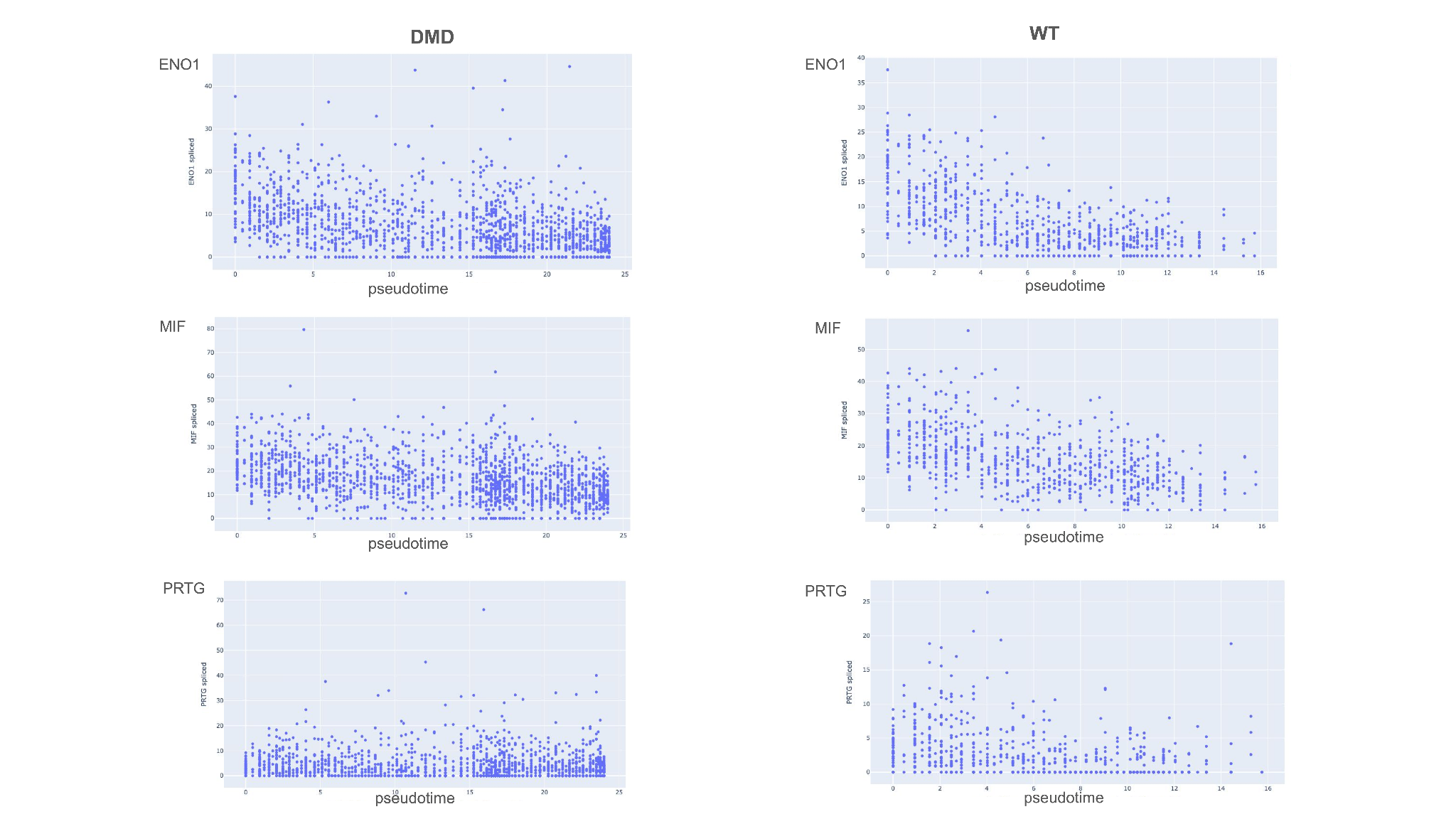

Figure S21: Waves of gene expression along the DMD and Healthy branches of the trajectory, for genes up-regulated on the DMD branch. The (pseudo)time scale corresponds to the rescaled latent time.

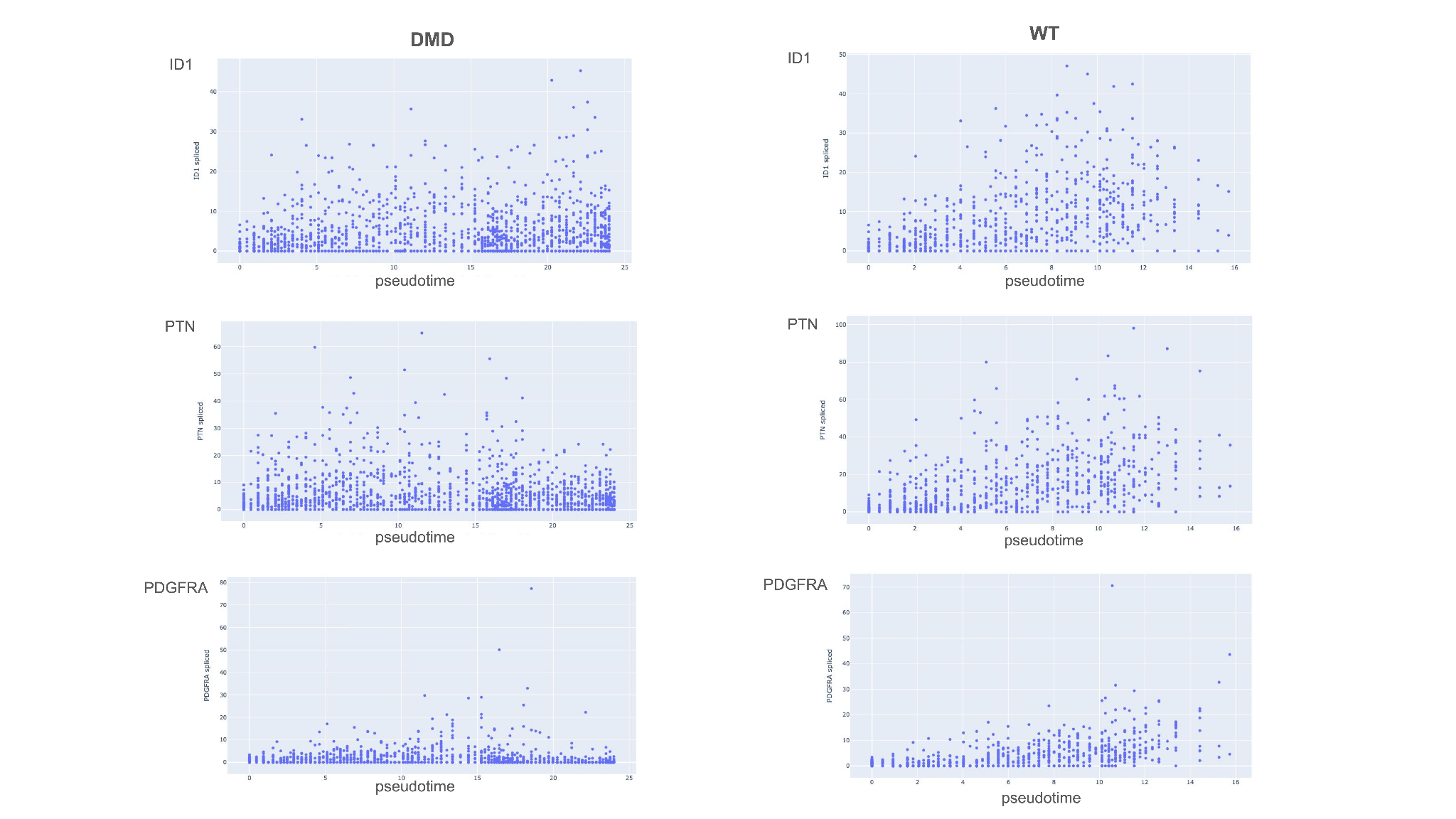

Figure S22: Waves of gene expression along the DMD and Healthy branches of the trajectory, for genes up-regulated on the Healthy branch. The (pseudo)time scale corresponds to the rescaled latent time.

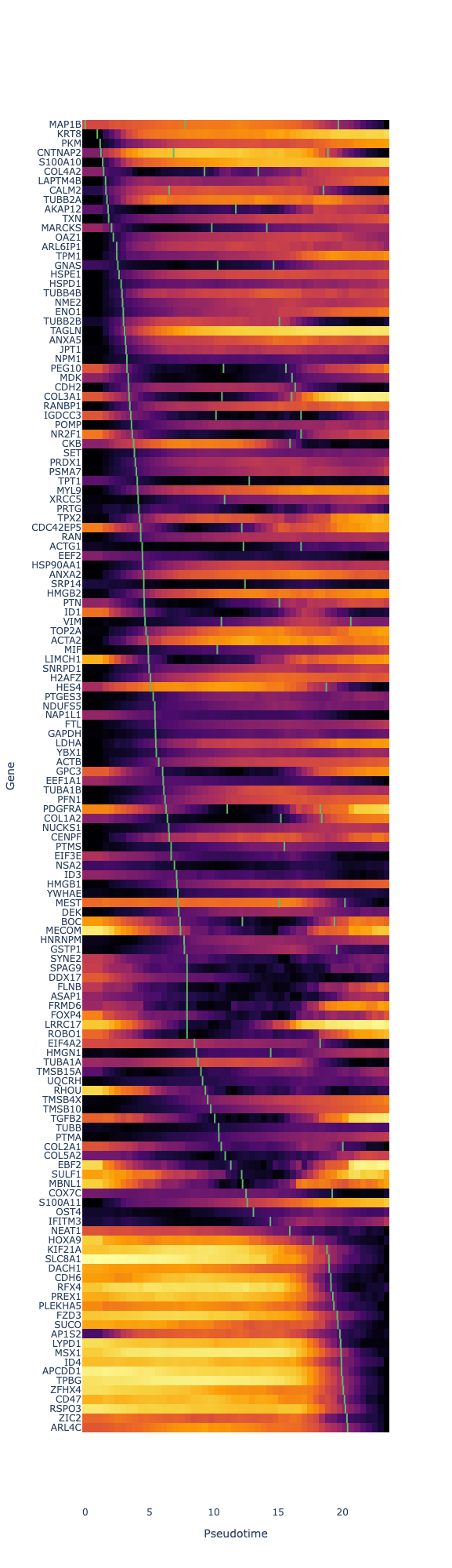

Figure S23: Gene expression heatmap focused on the 140 genes with the most significant wave for the DMD branch of the trajectory of interest. The green segments indicate waves half-(pseudo)time values. In this configuration, genes with early expression waves are located on top of the heatmap, and genes with later waves are located at the bottom.

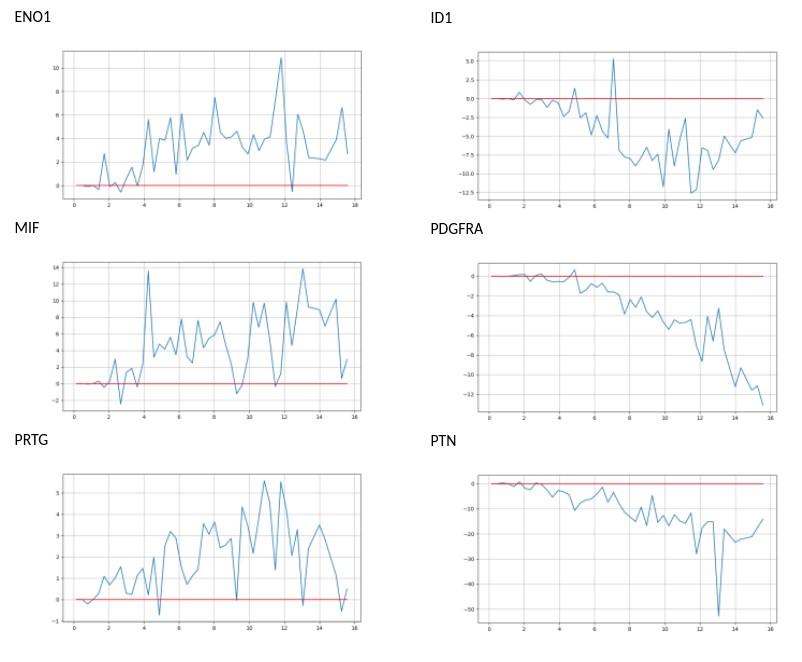

Figure S24: Difference of wave expression along pseudo time between the DMD and Healthy branches for the 6 toggle switch genes. A positive difference corresponds to a higher gene expression on the Healthy branch.

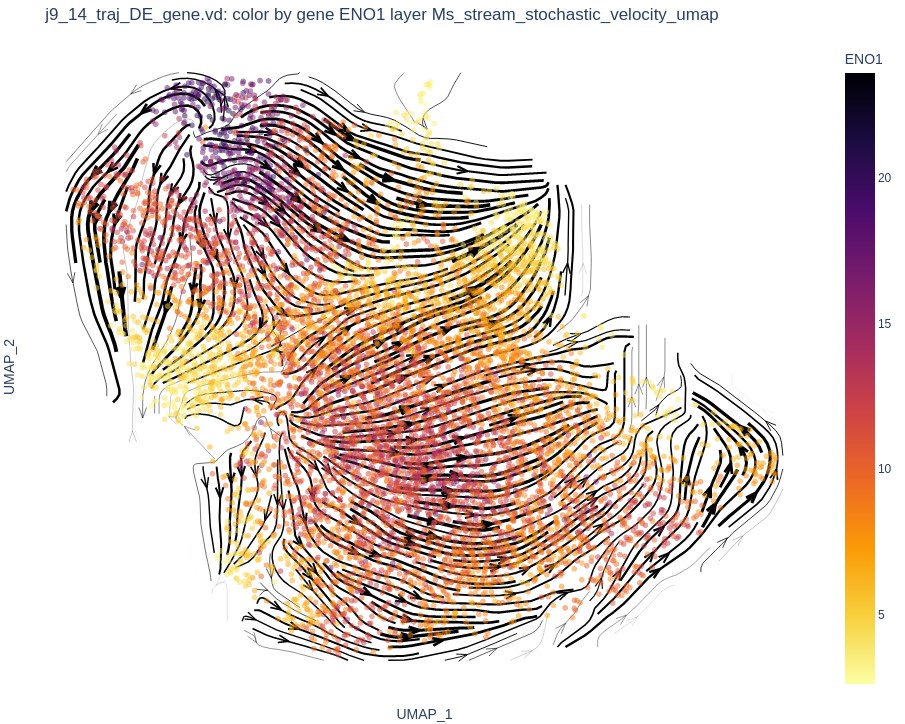

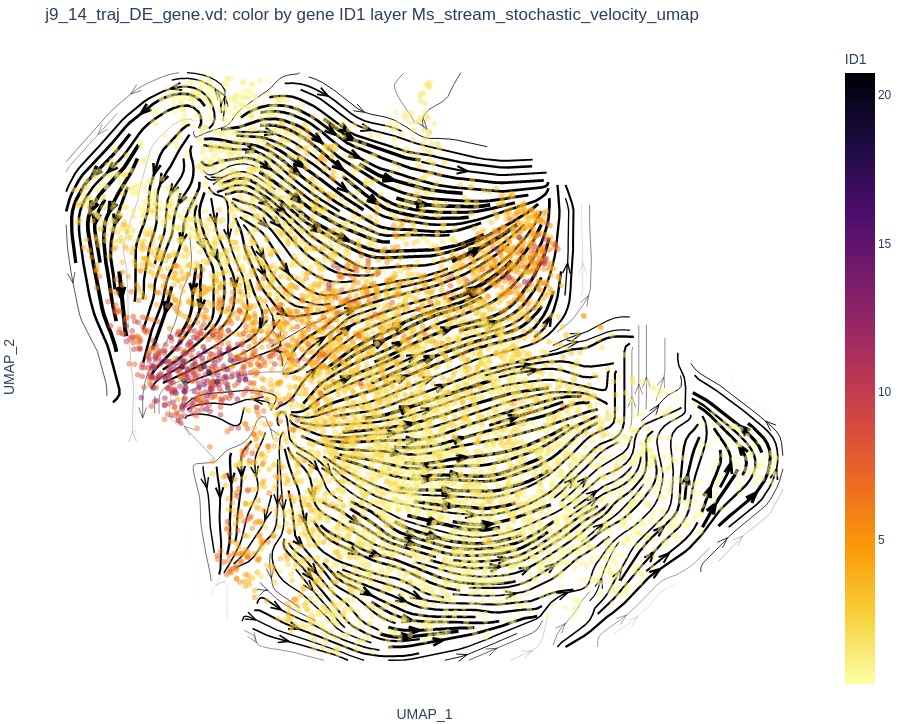

*ENO1*	*ID1*

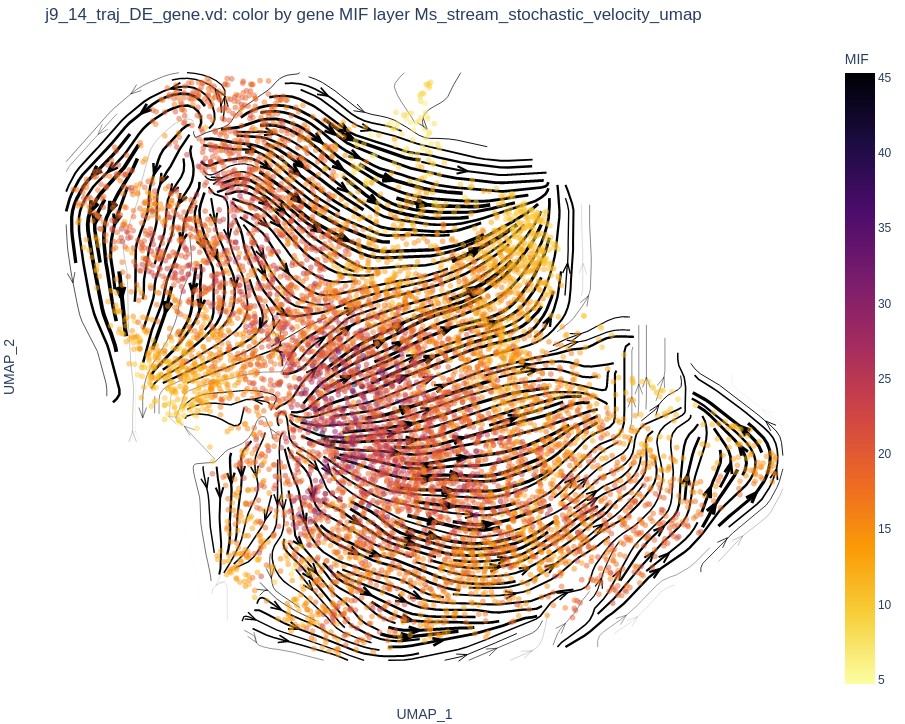

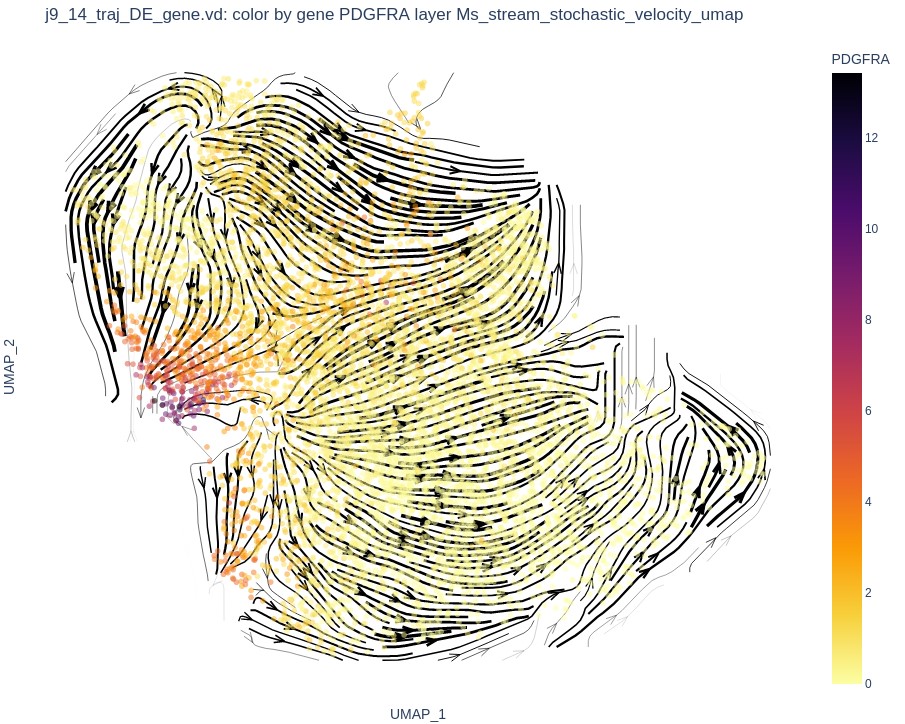

*MIF*	*PDGFRA*

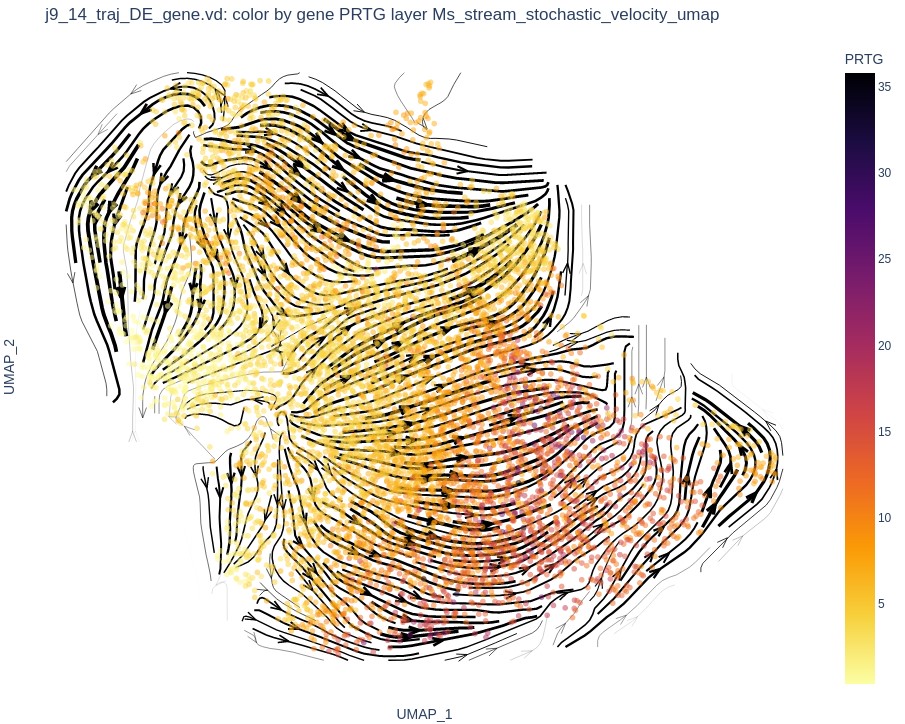

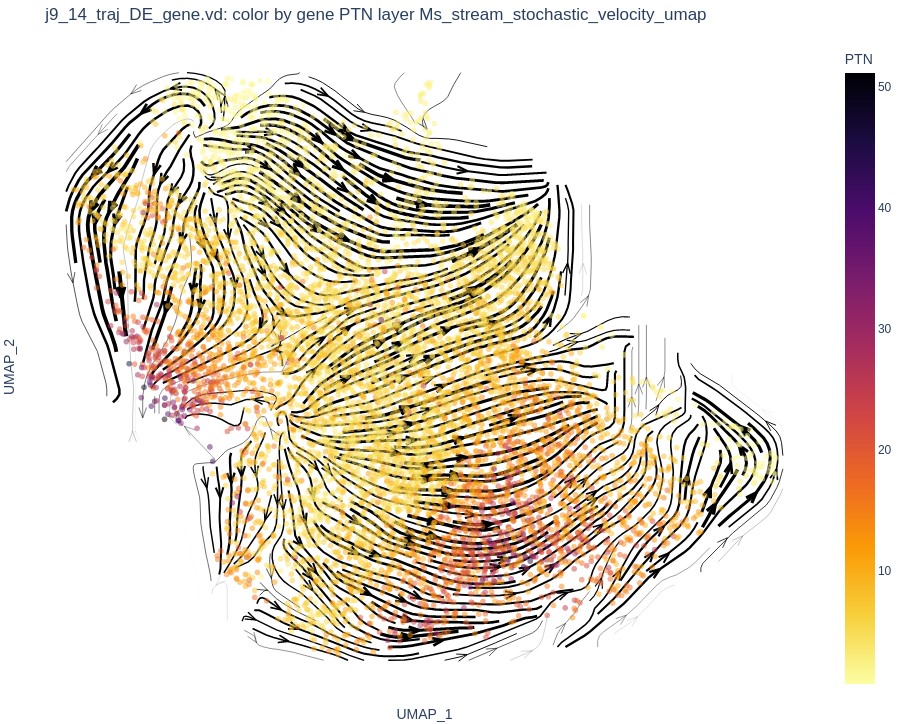

*PRTG*	*PTN*

Figure S25: Mean spliced layer for the 6 selected toggle switch genes

Healthy ENO1	DMD ENO1

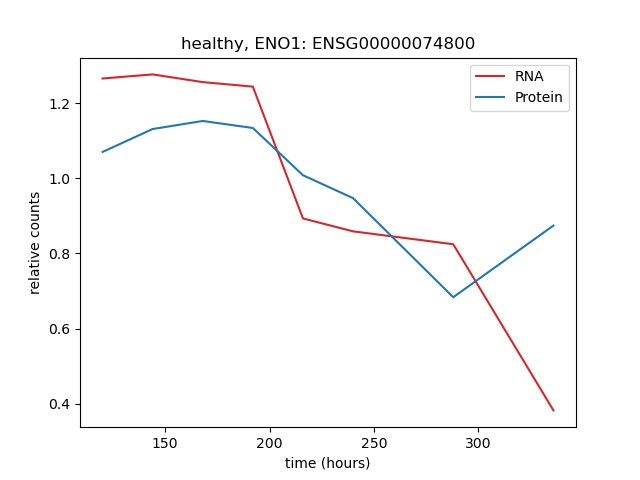

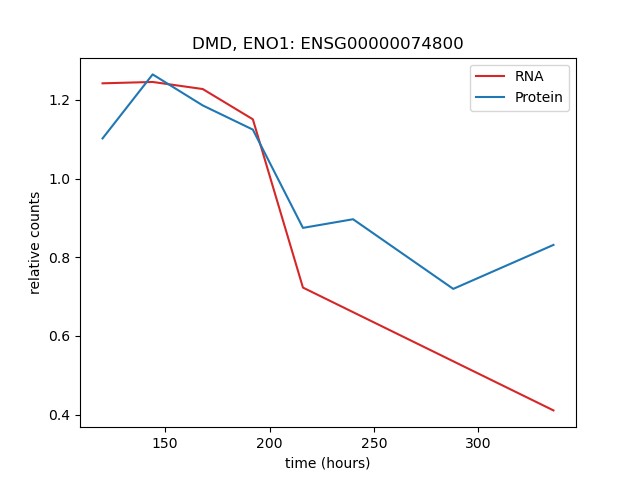

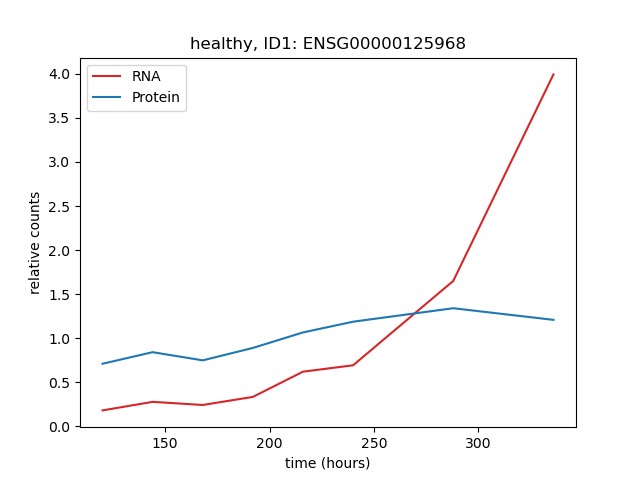

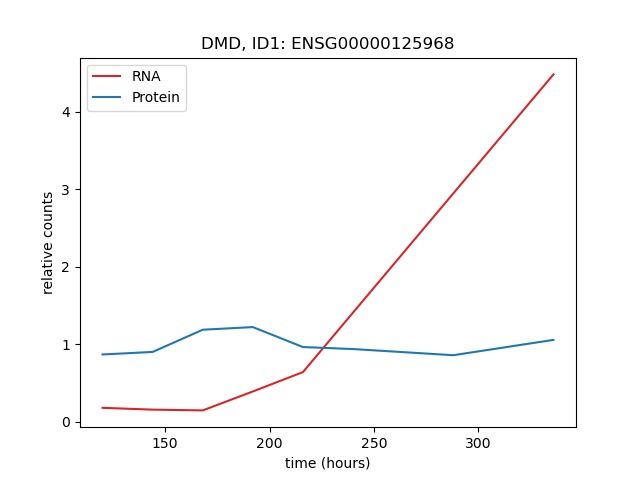

Healthy ID1	DMD ID1

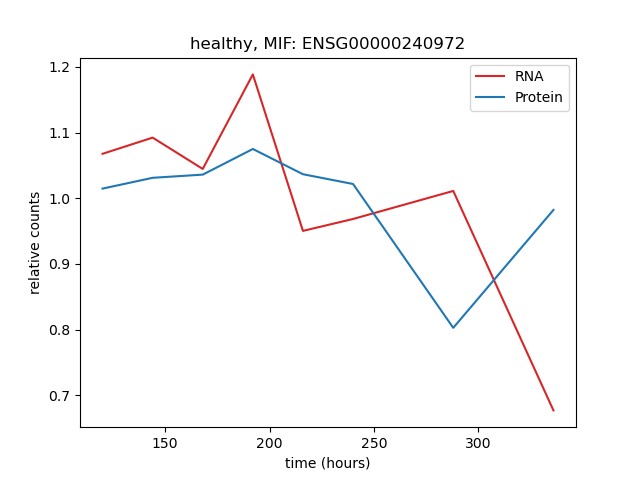

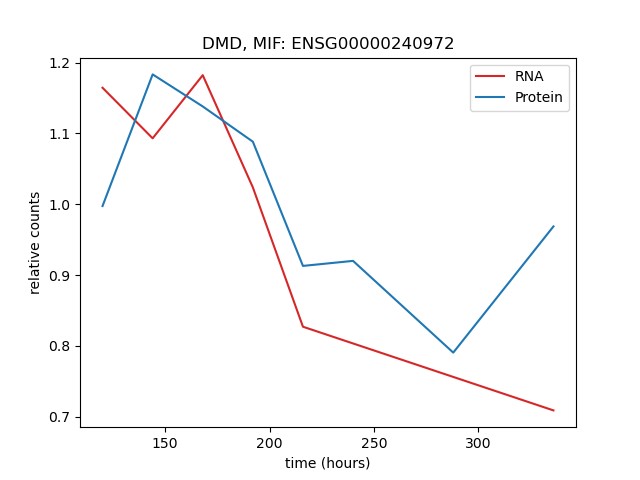

Healthy MIF	DMD MIF

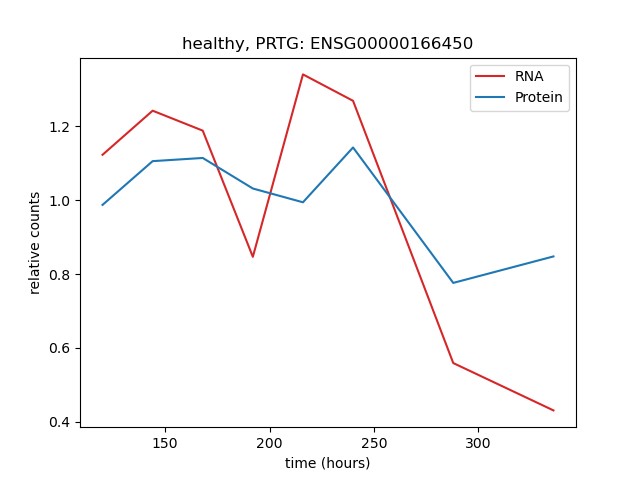

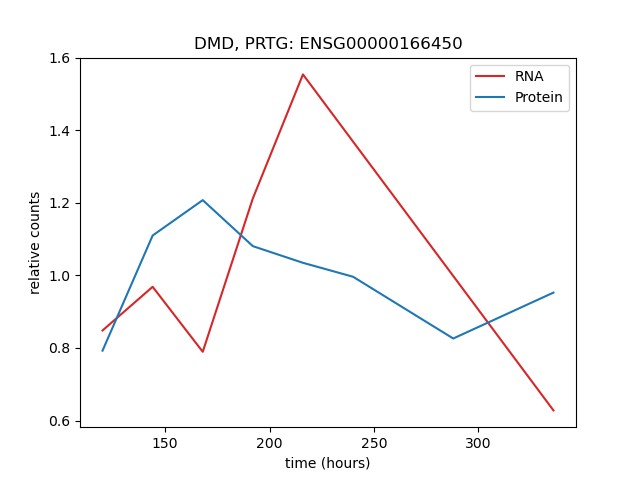

Healthy PRTG	DMD PRTG

Figure S26: Comparative evolution of protein and RNA levels for the 6 selected toggle switch genes on the Healthy and DMD branches. Note that PDGFRA and PTN proteins were not detected. For all genes and branches, the protein evolution is consistent with the RNA evolution, except for *ID1* where a de-correlation was observed at day 12 and day 14 (D12 and D14), particularly on the DMD branch.

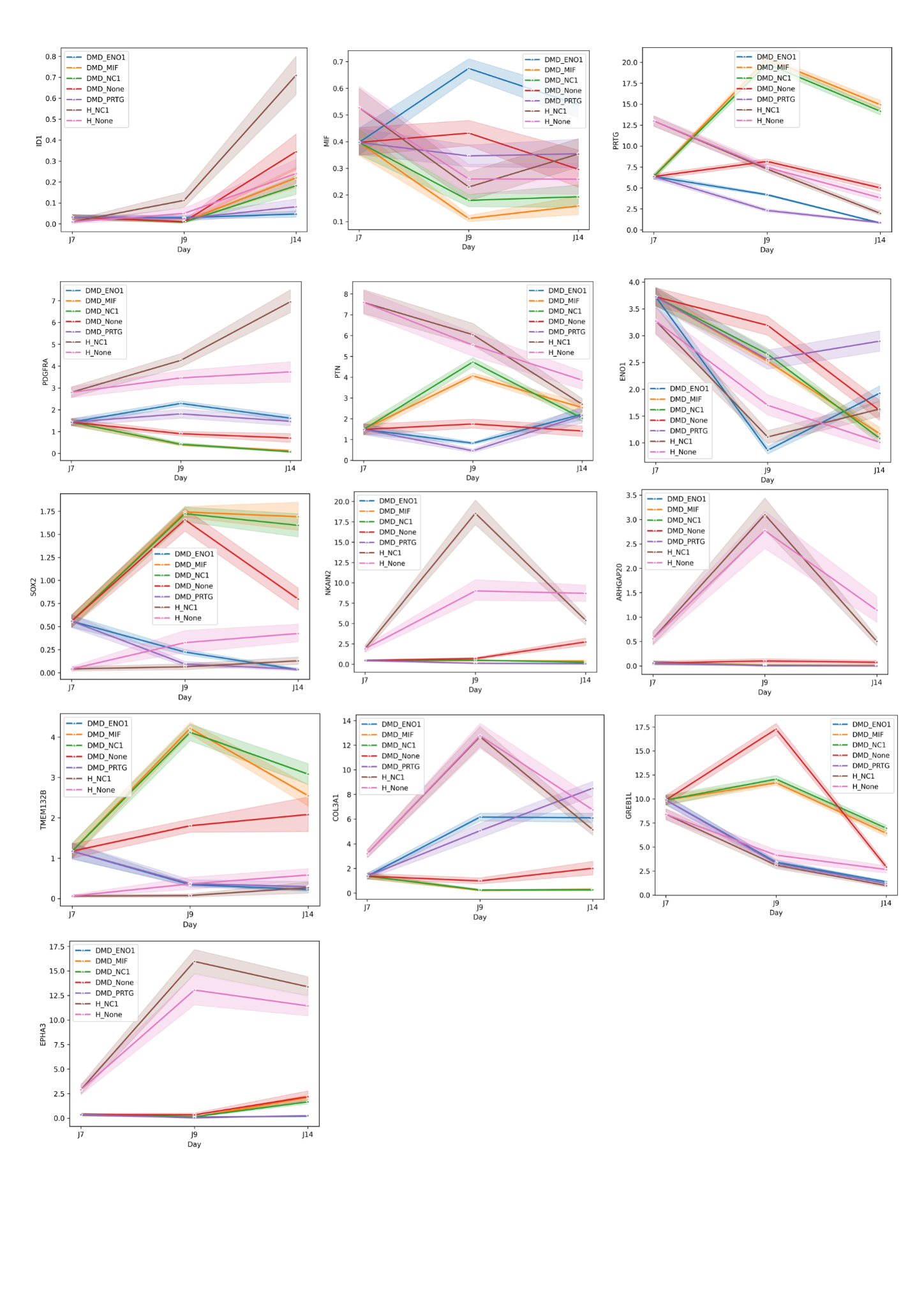

Figure S27: Mean and confidence interval of RNA counts vs time on scRNAseq2, colored by silencing RNA and genotype. All lines start at day 7 for the same genotype as the silencing RNA is only introduced at day 7.

| **Batch** | **Genotype** | **day** | **nb cell** |
| --- | --- | --- | --- |
| 1 | WT | 5 | 554 |
| 1 | WT | 6 | 306 |
| 1 | WT | 7 | 185 |
| 1 | WT | 8 | 344 |
| 1 | DMD | 5 | 355 |
| 1 | DMD | 6 | 691 |
| 1 | DMD | 7 | 262 |
| 1 | DMD | 8 | 618 |
| 2 | WT | 9 | 1306 |
| 2 | WT | 10 | 1275 |
| 2 | WT | 12 | 1452 |
| 2 | WT | 14 | 1297 |
| 2 | DMD | 9 | 1318 |
| 2 | DMD | 14 | 1002 |

Table S1: Description of the scRNAseq1 dataset

| **ensembl_id** | **gene_short_name** | **developmental_phase** |
| --- | --- | --- |
| ENSG00000204531.17 | POU5F1 | pluripotency |
| ENSG00000181449.3 | SOX2 | pluripotency |
| ENSG00000121570.12 | DPPA4 | pluripotency |
| ENSG00000111704.10 | NANOG | pluripotency |
| ENSG00000156574.9 | NODAL | pluripotency |
| ENSG00000147596.3 | PRDM14 | pluripotency |
| ENSG00000241186.9 | TDGF1 | pluripotency |
| ENSG00000125845.6 | BMP2 | mesoderm |
| ENSG00000165556.9 | CDX2 | mesoderm |
| ENSG00000163132.6 | MSX1 | mesoderm |
| ENSG00000164458.9 | T | mesoderm |
| ENSG00000149922.10 | TBX6 | mesoderm |
| ENSG00000124216.3 | SNAI1 | mesoderm |
| ENSG00000151379 | MSGN1 | mesoderm |
| ENSG00000106571.12 | GLI3 | somite |
| ENSG00000005102.12 | MEOX1 | somite |
| ENSG00000175745.11 | NR2F1 | somite |
| ENSG00000185551.14 | NR2F2 | somite |
| ENSG00000135903.18 | PAX3 | somite |
| ENSG00000105976 | MET | somite |
| ENSG00000125878 | TCF15 | somite |
| ENSG00000179111 | HES7 | somite |
| ENSG00000106003 | LFNG | somite |
| ENSG00000143632.14 | ACTA1 | muscle |
| ENSG00000175084.11 | DES | muscle |
| ENSG00000198947.15 | DMD | muscle |
| ENSG00000109063.14 | MYH3 | muscle |
| ENSG00000129152.3 | MYOD1 | muscle |
| ENSG00000122180.4 | MYOG | muscle |
| ENSG00000164093.15 | PITX2 | muscle |
| ENSG00000155657.26 | TTN | muscle |
| ENSG00000009709 | PAX7 | muscle |
| ENSG00000133020 | MYH8 | muscle |
| ENSG00000159251 | ACTC1 | muscle |

Table S2: Marker genes of muscle differentiation stage.

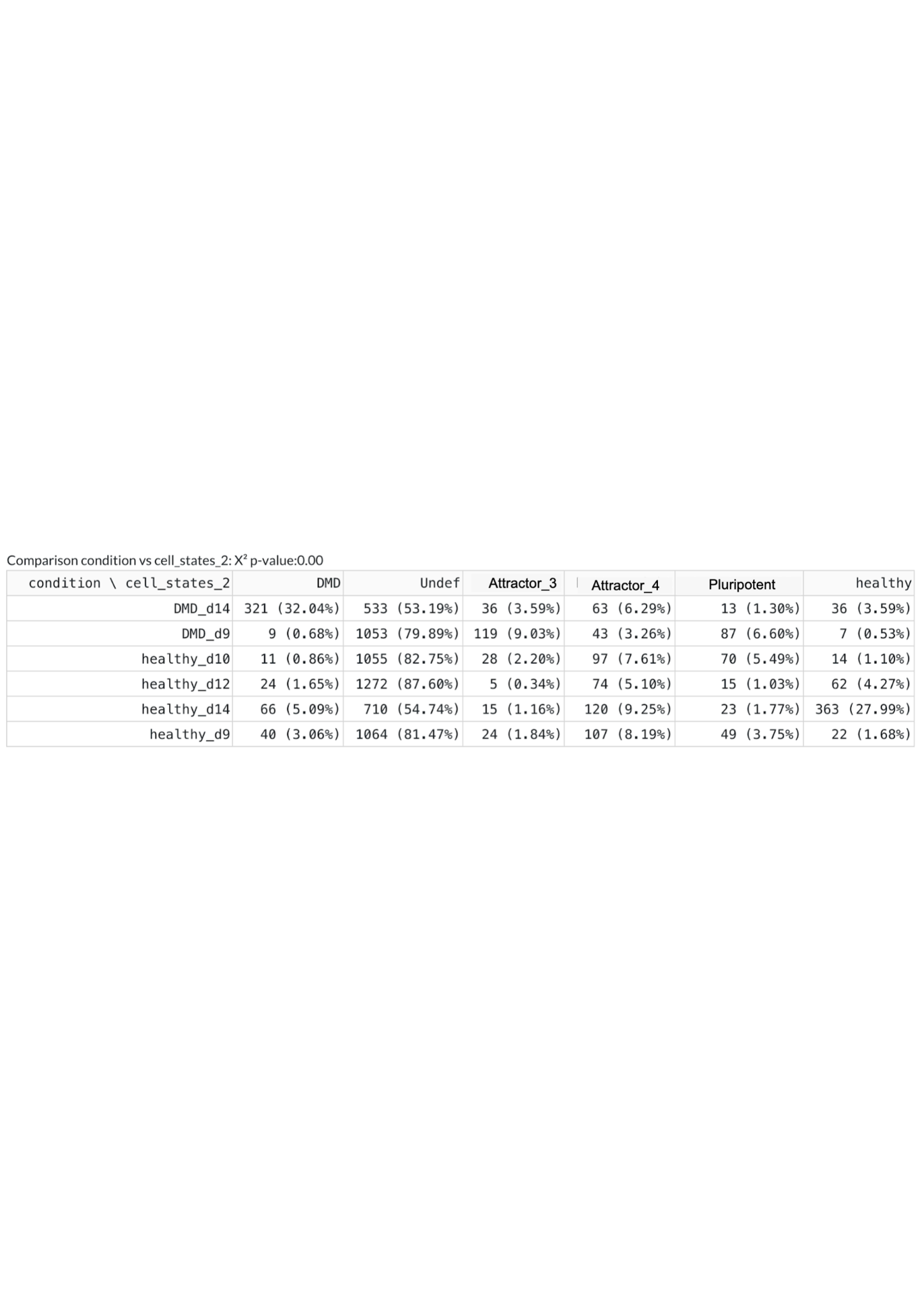

Table S3: Count table of the cells from each condition in each cluster, as defined in figure S10.

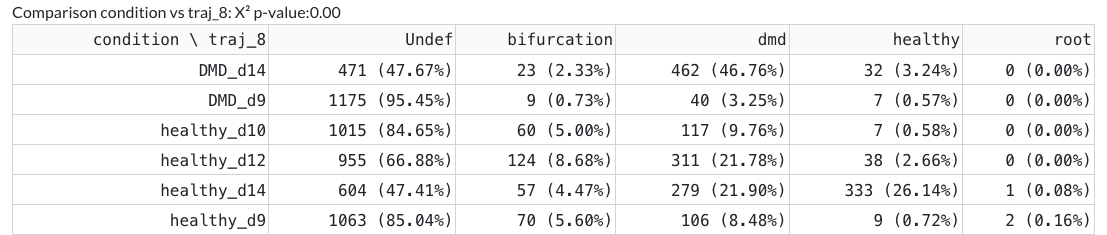

Table S4: Count table of the cells from each condition in each cluster along the trajectory, as defined in figure S17. Among the cell reaching a terminal state at day 14, 93.5% of mutant cells are in the DMD state DMD, 85.4% of WT cells are in Healthy state.

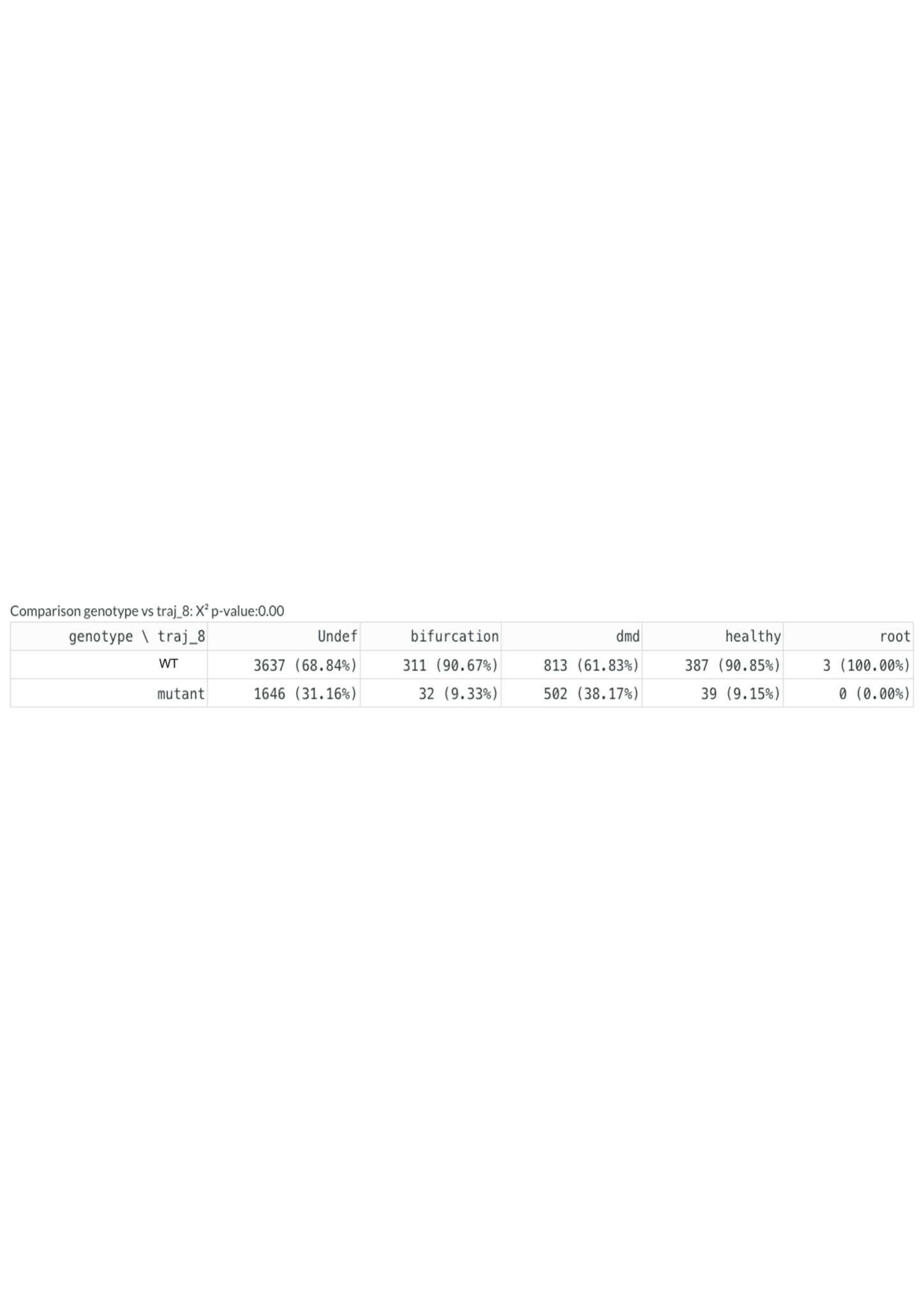

Table S5: Count table of the cells, by genotype, in each cluster along the trajectory as defined in figure S17. Among the cells in healthy final state, 90.85% are WT and 9.15% are mutant. Among the cells in the DMD final state, 38.17% are mutant and 61.83% are WT.
